# Catalytically distinct IDH1 mutants tune phenotype severity in tumor models

**DOI:** 10.1101/2024.04.22.590655

**Authors:** Mowaffaq Adam Ahmed Adam, Mikella Robinson, Ashley V. Schwartz, Grace Wells, An Hoang, Elene Albekioni, Cecilia Gallo, Grace Chao, Joi Weeks, Giovanni Quichocho, Uduak Z. George, Carrie D. House, Şevin Turcan, Christal D. Sohl

## Abstract

Mutations in isocitrate dehydrogenase 1 (IDH1) impart a neomorphic reaction that produces D-2-hydroxyglutarate (D2HG), which can inhibit DNA demethylases to drive tumorigenesis. Mutations affect residue R132 and display distinct catalytic profiles for D2HG production. We show that catalytic efficiency of D2HG production is greater in IDH1 R132Q than R132H mutants, and expression of R132Q in cellular and xenograft models leads to higher D2HG concentrations in cells, tumors, and sera compared to R132H. Though expression of IDH1 R132Q leads to hypermethylation in DNA damage pathways, DNA hypomethylation is more notable when compared to R132H expression. Transcriptome analysis shows increased expression of many pro-tumor pathways upon expression of IDH1 R132Q versus R132H, including transcripts of EGFR and PI3K signaling pathways. Thus, IDH1 mutants appear to modulate D2HG levels via altered catalysis, resulting in distinct epigenetic and transcriptomic consequences where higher D2HG levels appear to be associated with more aggressive tumors.

## Main

Isocitrate dehydrogenase 1 (IDH1) is a homodimeric enzyme that catalyzes the reversible NADP^+^-dependent oxidation of isocitrate (ICT) to α-ketoglutarate (αKG) in the cytoplasm and peroxisomes. Somatic mutations in IDH1 have been implicated in lower grade gliomas, chondrosarcomas, and intrahepatic cholangiocarcinomas^1^. These mutations typically ablate this conventional activity, but more importantly drive a neomorphic function: the NADPH-dependent conversion of αKG to the oncometabolite, D-2-hydroxyglutarate (D2HG)^2^. D2HG can competitively inhibit αKG-dependent enzymes such as the 5-methylcytosine hydroxylase TET2 and JmjC lysine demethylases to result in increased DNA and histone methylation, respectively^3–8^. We showed previously that mutations in IDH1 cause the CpG island methylator phenotype (CIMP)^9^, a clinical diagnostic and prognostic feature of tumors^10^.

Tumor-driving IDH1 mutations affect residue R132, with R132H and R132C the most common, though more rare mutations, including R132G, R132L, R132S have been reported^2,11,12^. Residue 132 interacts with the C3 carboxylate of ICT^2,13–15^ and stabilizes an important regulatory domain, the α10 helix^2,13,14,16,17^. Thus, loss of arginine at residue 132 helps favor a closed conformation to drive the equilibrium towards αKG and NADPH binding, explaining why there is some permissiveness in mutations^2,15^. There has been interest in determining if different IDH1 mutations lead to different features in IDH1-driven tumors. Glioma tissue (expressing IDH1 R132H/C/G mutations) and cell lines (expressing IDH1 R132H/C/G/S/L mutations) showed varying concentrations of D2HG depending on the IDH1 mutation present, with R132H associated with the lowest levels of D2HG^18,19^. We previously reported that purified IDH1 R132Q was far more catalytically efficient at D2HG production than R132H^20,21^ due to activating structural features^22^. An extensive study on astrocytomas that were either R132H-mutated or non-R132H IDH1/2-mutated showed that non-R132H IDH1/2-mutated tumors had overall increased DNA methylation, decreased gene expression, and better prognosis, suggesting an intriguing possibility of differences in methylation phenotype intensity within IDH1 mutants and/or among IDH1 and IDH2 mutants^23^. Together, these studies highlight an important unresolved question: do different concentrations of D2HG resulting from different IDH1 mutations lead to changes in tumor phenotype or phenotype intensity? If increased catalytic rates conferred among some IDH1 mutants can indeed lead to higher D2HG concentrations in tumor models, understanding the impact of these mutations on tumor growth rates, epigenetic, and transcriptomic profiles can ultimately help us better predict patient prognosis.

Here, we sought to determine the *in vitro* and *in vivo* consequences of two IDH1 mutants with profoundly different kinetic parameters. We generated ectopic mouse tumor xenografts from either glioma or sarcoma cell lines we modified to stably over-express IDH1 WT, R132H or R132Q and performed comprehensive epigenomic and transcriptomic analyses. We found that tumors expressing R132Q had increased D2HG levels and larger tumors compared to WT and R132H, but the glioma and chondrosarcoma xenograft tumor models showed highly variable consequences on genome methylation and gene expression, with increased pro-tumorigenic pathways upregulated upon expression of IDH1 R132Q.

## Results

### D2HG levels in IDH1 R132Q tumor models are higher

Though we have shown that IDH1 R132Q mutant homodimers produce D2HG more efficiently than R132H mutant homodimers^20–22^, here we made 1:1 mixtures of purified WT and mutant IDH1 enzymes to mimic the heterozygosity in patients. These IDH1 WT:R132Q mixtures, which may form heterodimers, retained superior catalytic efficiency compared to WT:R132H, with 9.5-fold more efficiency for D2HG production (Supplementary Fig. 1). This was driven both by an increase in *k*_cat_ and decrease in *K*_m_.

Though catalytic parameters identify intrinsic enzyme properties, this ignores the complexities of the cellular environment. As patient-derived IDH1 R132Q-driven tumor cells were unavailable due to the rarity of this mutation, we generated isogenic cell lines stably overexpressing IDH1 WT, R132H, R132Q, or empty vector. We selected well-characterized cancer-derived and non-cancer-derived cell lines commonly used to model IDH1 mutations in gliomas and chondrosarcomas: U87MG glioma cells, normal human astrocytes (NHA), C28 chondrocytes, and a modified HT1080 chondrosarcoma line with the endogenous IDH1 R132C mutation removed to generate IDH1^-/+^, designated as HT1080*. Interestingly, we were unable to generate clones that sustained high expression of IDH1 R132Q (Supplementary Fig. 2). Despite consistently lower expression of IDH1 R132Q relative to R132H, we nonetheless found that D2HG concentrations were significantly higher in this more catalytically active mutant (Fig. 1a) in both cancerous (U87MG, HT1080*) and non-cancerous cell lines (NHA, C28).

**Fig. 1.**
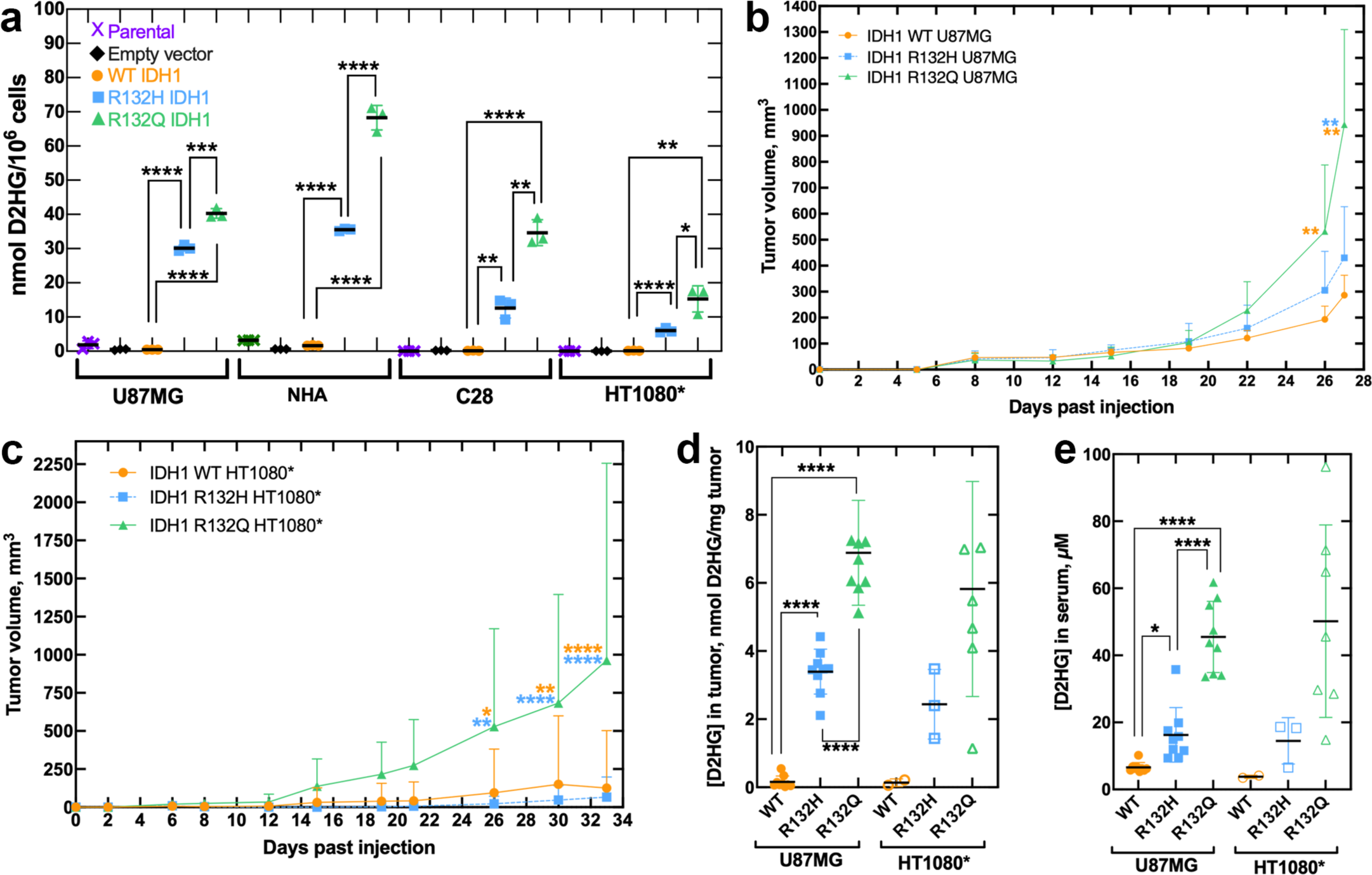
Catalytic efficiency of IDH1 mutants drive D2HG levels in biochemical, cellular, and *in vivo* models. **a**, Parental (purple x’s), empty vector (EV, black diamonds), IDH1 WT (orange circles), IDH1 R132H (blue squares), or IDH1 R132Q (green triangles) was stably overexpressed in glioma cells (U87MG), normal human astrocytes (NHA), chondrocytes (C28), and fibrosarcoma cells where an endogenous IDH1 R132C mutation was removed (i.e. HT1080^+/-^, denoted HT1080*), and cellular D2HG levels were quantified in three biological replicates per cell type. Significant differences involving parental and empty vector (EV) are not highlighted to improve clarity. **b**, U87MG and **c**, HT1080* lines stably overexpressing IDH1 WT, R132H, or R132Q were used to generate mouse xenografts and tumor volume was measured over time. All 27 mice (nine per sample group) formed U87MG xenograft tumors, and the mean of the nine biological replicates with standard deviation (S.D.) is shown in (b). Only two, three, or seven mice formed HT1080* xenograft tumors upon IDH1 WT, R132H, or R132Q expression, respectively, and the mean of these biological replicates with S.D. is shown in (c). In (b, c), significant differences from IDH1 R132Q are indicated with *p* value stars colored according to the corresponding comparison group (WT in orange, R132H in blue). **d**, Tumors, and **e**, sera were harvested from the mice described in (b) and (c), and D2HG was quantified. In all panels (a-e), *p* values were determined by unpaired t tests (a) or ordinary one-way ANOVA (b-e), with **** *p* ≤ 0.0001, *** *p* ≤ 0.001, ** *p* ≤ 0.01, * *p* ≤ 0.05 for pairwise comparisons between IDH1 WT, R132H, and R132Q.

We next sought to understand the consequences of mutants with differing catalytic efficiency in *in vivo* models of mutant IDH1 tumors. To increase the likelihood of tumor formation, we used IDH1 WT-, R132H-, or R132Q-expressing glioma (U87MG) and fibrosarcoma (HT1080*) cells to generate subcutaneous mouse xenografts. For the U87MG series, tumors formed in 100% of the mice (27/27, 9 per group), with no significant differences in days to tumor initiation or tumor density (Supplementary Fig. 3). However, IDH1 R132Q U87MG tumors had significantly higher volume (Fig. 1b), mass, and growth rate (Supplementary Fig. 3) compared to WT and R132H. Tumors were formed in 22% (2/9), 30% (3/10), and 78% (7/9) of the IDH1 WT-, R132H-, or R132Q-expressing HT1080* mouse xenografts, respectively. Here, tumor volume was larger (Fig. 1c), and tumor initiation occurred significantly earlier for IDH1 R132Q mutants compared to WT and R132H, though tumor mass and tumor density showed no significant difference (Supplementary Fig. 4).

We harvested all available tumors and sera from mice and found that D2HG levels were significantly higher in IDH1 R132Q-expressing U87MG xenografts relative to WT and R132H (Fig. 1d, 1e), even though expression of R132H and WT was higher compared to R132Q (Supplementary Fig. 5, Supplementary Table 1). These trends were recapitulated in the HT1080* xenografts, though significance was not reached (Fig. 1d, 1e). There were few obvious trends among D2HG concentrations and tumor features within biological replicates, though modest correlation in serum D2HG concentration and tumor mass as a percentage of body mass (Supplementary Fig. 6) was observed. Together, these tumor models showed that expression of IDH1 R132Q was associated with more catalytically efficient D2HG production, higher cellular, tumor, and serum levels of D2HG, and larger tumors compared to R132H.

### Epigenomic and transcriptomic features of mutant IDH1 chondrosarcoma models

To understand the mechanism(s) behind the differences in tumor size in the IDH1 R132Q versus R132H and WT tumors, we used reduced representation bisulfite sequencing (RRBS) to analyze genome-wide DNA methylation in representative xenograft tumors. We first performed clustering analysis for the HT1080* xenografts based on the degree of CpG methylation of the most variable probes with the highest S.D. (Fig. 2a). There was modest clustering based on genotype, with HT1080* xenografts expressing IDH1 WT having the most variation. This is perhaps unsurprising given the wide span in tumor weight and low *n* for WT IDH1-expressing HT1080* tumors (Supplementary Table 1). Better separation of the two mutants was achieved via PCA that included D2HG tumor concentrations (Fig. 2b). When comparing distribution of methylation values for all CpG sites, there was a modest shift in hypermethylation in mutant versus WT HT1080* xenografts (Fig. 2c). We then analyzed the methylation levels of CpG sites that correlated with tumor D2HG amounts in HT1080* xenografts and identified the loci that were positively correlated with D2HG (Fig. 2d). Next, we identified the pairwise differentially methylated loci among all three genotypes in HT1080* tumors and found IDH1 R132Q induced hypermethylation in a larger number of CpG sites than R132H when compared to WT (Fig. 2e). In general, both IDH1 R132H and R132Q chondrosarcoma tumor models showed the same hypermethylation localization trends compared to WT, with promoter and intronic regions showing the most hypermethylation (Fig. 2f). However, when comparing the two mutants, significantly increased hypomethylation was observed upon R132Q expression versus R132H (Fig. 2g-i, Supplementary Fig. 7), especially in promoter regions and CpG islands (Fig. 2f, Supplementary Fig. 8).

**Fig. 2.**
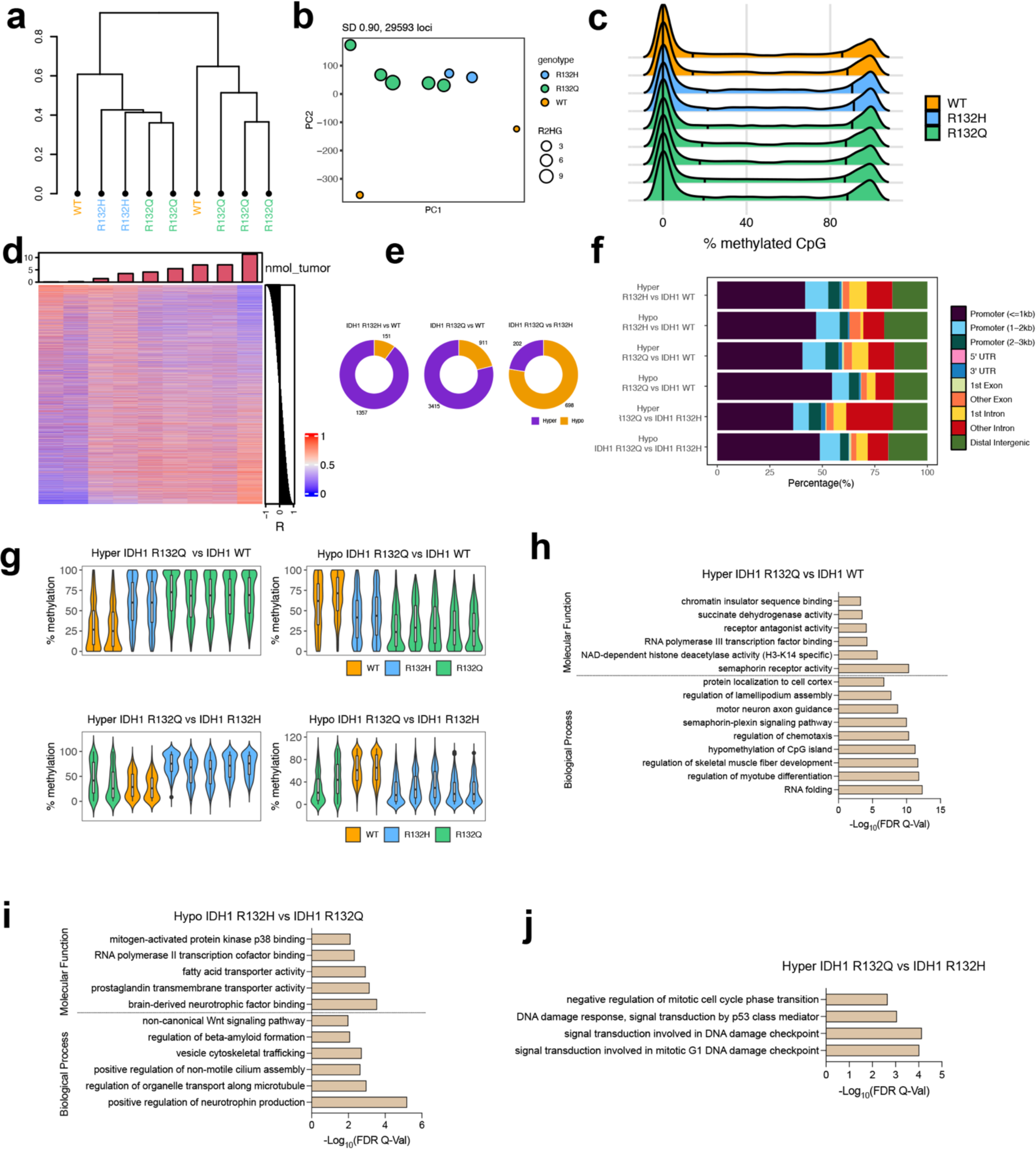
IDH1 R132Q-induced methylation remodeling differs from IDH1 R132H in HT1080* tumors. Two of two, three of three, and five of seven (randomized selection) of the xenograft tumors that formed HT1080* xenograft tumors upon IDH1 WT, R132H, or R132Q expression, respectively, were evaluated as biological replicates by RRBS, though one of the R132H xenograft tumors failed analysis. **a**, Hierarchical clustering of the most variable CpG sites (S.D. > 0.90, n = 29,593) in the HT1080* RRBS cohort. **b**, PCA plot of the most variable CpG sites (standard deviation > 0.90) in the HT1080* RRBS cohort. The size of each dot corresponds to the level of D2HG (nmol tumor) measured in each tumor sample. Colors indicate genotype. **c**, Global methylation profiles of the HT1080* RRBS tumor cohort showing percent (%) methylation values of CpG sites for each sample. Black vertical lines indicate quantiles, colors indicate genotype. **d**, Heatmap showing the correlation between D2HG (nmol tumor) and methylation values of the top 1% of the most variable CpG sites. Pearson correlation coefficients (R) are shown on the right of the heatmap. The amount of D2HG in tumors is depicted as a bar plot above the heatmap. **e**, Number of differentially methylated CpG sites (DMS, % methylation difference ≥ 20 and q-value < 0.05) are shown. Left, IDH1 R132H vs IDH1 WT; middle, IDH1 R132Q vs IDH1 WT; right, IDH1 R132Q vs IDH1 R132H. hyper, hypermethylated; hypo, hypomethylated. **f**, Violin plots showing the distribution of percent (%) methylation of hypermethylated loci in IDH1 R132Q vs IDH1 WT (top left), hypomethylated loci in IDH1 R132Q vs IDH1 WT (top right), hypermethylated loci in IDH1 R132Q vs IDH1 R132Q (bottom left), and hypomethylated loci in IDH1 R132Q vs IDH1 R132H (bottom right). For clarity, samples not included in the pairwise statistical comparisons are also plotted. **g**, Distribution of the genomic features for the hyper- and hypo-methylated loci for all comparisons. **h-j**, Gene ontology enrichment (using GREAT toolbox) of the hypermethylated (hyper) loci in IDH1 R132Q compared to **h,** IDH1 WT, **i**, hypomethylated (hypo) loci in IDH1 R132Q vs IDH1 R132H, and **j**, hypermethylated loci in IDH1 R132Q vs IDH1 R132H samples.

To compare the expression patterns of genes in the WT-, R132H-, and R132Q-driven xenografts, we also performed RNA-seq analysis. Consistent with both mutants being hypermethylated within the promoter regions relative to WT, which would predict increased gene silencing, there were more down-regulated than up-regulated genes for R132H and R132Q HT1080* xenografts relative to WT (Supplementary Table 2). Similarly, consistent with an increase in hypomethylation of CpG sites in R132Q versus R123H, which would predict an increase in gene expression, there were more upregulated genes in R132Q than downregulated compared to R132H (Supplementary Table 2). Together, these data suggest that in chondrosarcoma tumor models, expression of IDH1 R132 mutants led to increased hypermethylation compared to WT, while R132Q was associated with increased regions of hypomethylation compared to R132H.

### Pathways hypermethylated and down-regulated in IDH1 R132Q chondrosarcoma models

Hypermethylated pathways in IDH1 R132Q-versus R132H-expressing chondrosarcoma models included cell cycle, DNA damage response, and DNA damage checkpoint signal transduction (Fig. 2h), suggesting the DNA damage response is likely inhibited. Notably, IDH1 R132Q had higher methylation at the promoter of mitotic arrest deficient 2-like protein 2 (*MAD2L2*, also known as *REV7*) relative to R132H. Consistent with this increased methylation, RNAseq analysis showed expression of *MAD2L2* trended lower in R132Q-expressing HT1080* tumors compared to R132H (*p* = 0.028, *p*_adj_ = 0.21). MAD2L2 plays a role in translesion DNA synthesis and double stranded DNA break repair, promoting progression of stalled replication forks past lesions and driving DNA repair pathways^24,25^. Loss of MAD2L2 leads to genomic damage as a result of stalled replication forks^26^ and increased proliferation and migration^27^, and thus has been posited to be a tumor suppressor though it may have an oncogenic role in some contexts^28^. Significant hypermethylation within the promoter of growth arrest and DNA-damage-inducible protein γ (*GADD45G*) occurred in IDH1 R132Q versus R132H HT1080* xenograft tumors, though this gene did not appear in our RNAseq analysis. GADD45G is a tumor suppressor that is involved in a host of cellular processes, including DNA repair^29^, cell cycle arrest^30^, and apoptosis^31^. Notably, its silencing or downregulation has been implicated in AML^32^ and breast cancer^33^. GADD45G promotes cell differentiation through negative regulation of the phosphoinositide 3-kinase/AKT/mTOR (PI3K/AKT/mTOR) pathway, and, consequently, its downregulation leads to activation of this pathway to drive cancer^34^.

Many pathways showed significant hypermethylation in IDH1 R132Q vs. WT HT1080* xenografts, including pathways involved in altered transcription and cell movement (Fig. 2i). Pathways associated with morphogenesis and differentiation were also hypermethylated in R132Q relative to WT, including significant hypermethylation of Wnt pathway members *WNT9A*, *WNT10A*, frizzled-9 (*FZD9*), palmitoleoyl-protein carboxylesterase (*NOTUM*), T-cell specific transcription factor 7 (*TCF7*), and *NKD1*. Indeed, RNAseq analysis showed that *WNT9A* was significantly down-regulated in R132Q versus WT HT1080* xenografts, with *FZD9* and *TCF7* transcripts trending down but not achieving significance when adjusted for multiple comparisons (Supplementary Table 3).

RNAseq analysis also identified transcript downregulation not necessarily driven by genome hypermethylation. For example, transcripts involved in collagen/extracellular matrix (ECM) pathways were downregulated upon R132Q expression compared to both R132H and WT HT1080* xenografts (Fig. 3, Supplementary Fig. 9). This included decreased *COL9A2* transcripts upon expression of R132Q versus WT, a type IX collagen often downregulated in high-grade versus low-grade chondrosarcomas^35^ (Supplementary Table 3). Annexin-A6 (*ANXA6*) transcripts were also downregulated in R132Q compared to both R132H and WT chondrosarcoma models (Supplementary Table 3). ANXA6 mediates cholesterol transport and participates in endocytosis and exocytosis^36^, and can serve as a tumor suppressor due to its ability to negatively regulate EGFR phosphorylation^37^. Supportive of removing a negative regulator, *EGFR* expression was higher in R132Q versus R132H and WT HT1080* xenografts (Supplementary Table 3). Lipid catabolism/oxidation pathways were also downregulated in R132Q versus R132H (Fig. 3), suggesting that lipids as a fuel source is more critical in R132H than R132Q; indeed, β-oxidation has been shown to be an important adaptive mechanism in R132H tumor models^38^. RNAseq analysis also highlighted that cell surface projection pathways were more important in R132H-expressing HT1080* xenografts compared to R132Q or WT (Supplementary Fig. 10). Overall, compared to IDH1 R132H, R132Q-expressing chondrosarcoma tumor models experienced significant changes in pathways involving DNA damage, Wnt and EGFR signaling, and collagen/ECM remodeling that was facilitated by altered CpG methylation and non-CpG-methylation pathways.

**Fig. 3.**
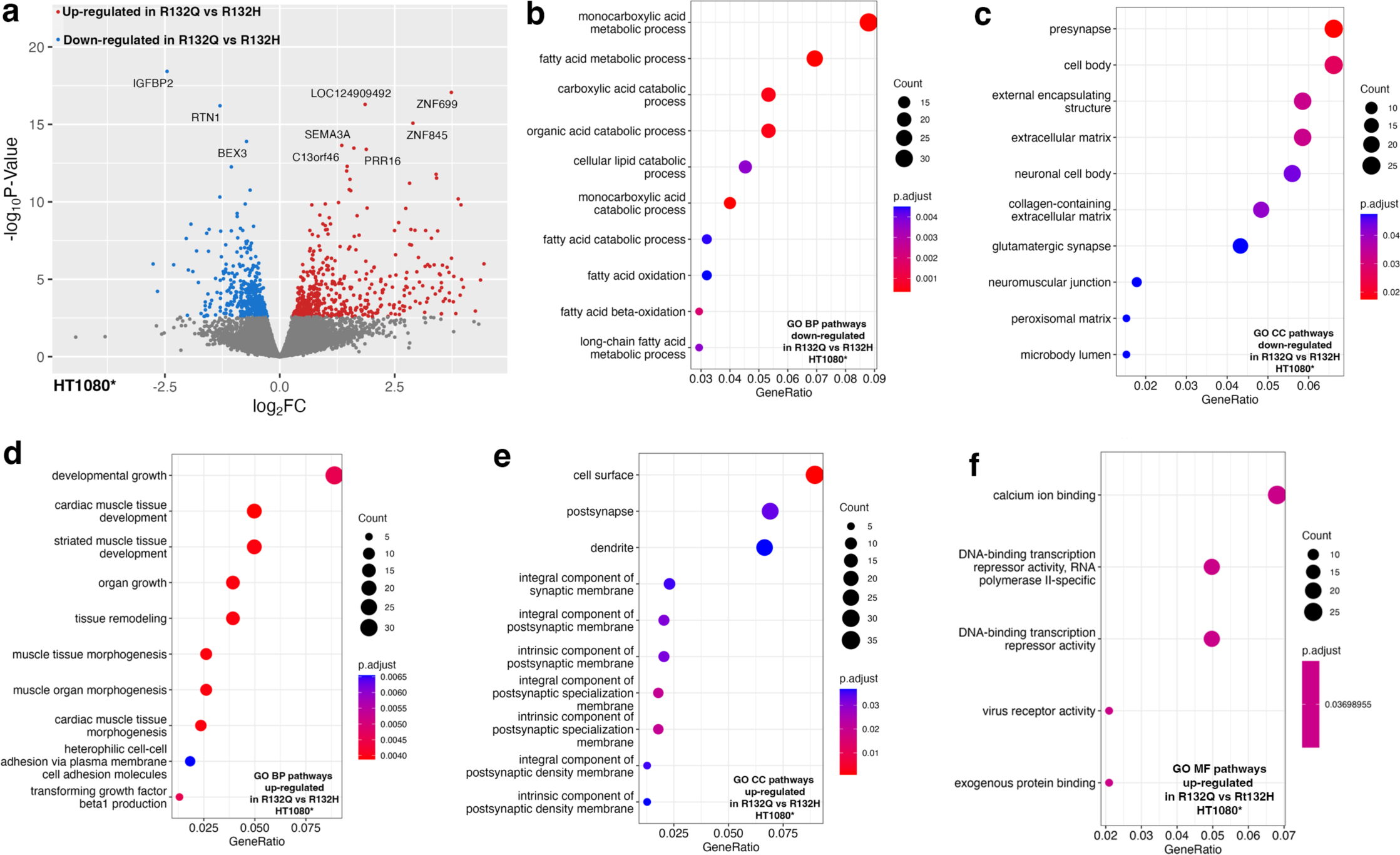
Transcriptome analysis using RNAseq of HT1080* xenograft tumors comparing IDH1 R132Q and IDH1 R132H. Two of two, three of three, and five of seven (randomized selection) of the xenograft tumors that formed HT1080* xenograft tumors upon IDH1 WT, R132H, or R132Q expression, respectively, were evaluated as biological replicates by RNAseq. Down-regulated pathways refer to categories where most genes show decreased expression levels in R132Q versus R132H IDH1. Up-regulated pathways refer to categories where most genes show increased expression levels in R132Q versus R132H IDH1. Number of genes is indicated in the count, with the *p*_adjusted_ value indicated by color. **a**, Volcano plot of differentially expressed transcripts comparing expression of IDH1 R132Q versus IDH1 R132H. **b**, Biological pathways (BP) down-regulated in IDH1 R132Q versus IDH1 R132H. **c**, Cellular component (CC) pathways down-regulated in IDH1 R132Q versus IDH1 R132H. **d**, BP up-regulated in IDH1 R132Q versus IDH1 R132H. **e**, CC pathways up-regulated in IDH1 R132Q versus IDH1 R132H. **f**, Molecular function (MF) pathways up-regulated in IDH1 R132Q versus IDH1 R132H.

### Pathways hypomethylated and up-regulated in IDH1 R132Q chondrosarcoma models

As hypomethylated CpG regions was an important difference between R132Q and R132H HT1080* xenografts, we also assessed the pathways affected by decreased methylation, and thus likely upregulated. Pathways predicted to be hypomethylated in R132Q versus R132H chondrosarcoma models involved p38 MAPK, which helps regulate cell differentiation, growth, and death^39^, RNA pol II transcription cofactor binding, vesicle trafficking, fatty acid and organelle transport pathways, and non-canonical Wnt signaling that can drive proliferation and migration in cancer^40^ (Fig. 2j). Thus, Wnt signaling alterations resulted from both hypermethylation and hypomethylation in IDH1 R132Q tumors.

Similarly, GO pathways upregulated based on increased transcripts in IDH1 R132Q versus R132H HT1080* xenografts included pathways associated with growth, remodeling, morphogenesis, and transcription repression (Fig. 3), but there was minimal overlap of genes significantly hypomethylated with elevated transcripts. For example, the *MRAS* promoter was hypomethylated in R132Q-expressing xenografts compared to R132H, though this did not translate to a significant increase in *MRAS* expression. To understand the consequences of a less methylated genome in R132Q than R132H, we also compared pathways with hypomethylation and increased transcripts in IDH1 R132Q versus WT. We observed hypomethylation of pathways including vesicle loading of neurotransmitters, and indeed RNAseq analysis highlighted an increase in pathways associated with endocytic vesicles, and neuronal projection and axon development/guidance (Supplementary Fig. 9)

RNAseq analysis again showed many cancer pathways upregulated in R132Q HT1080* xenografts versus R132H outside of genes identified in RRBS analysis, including proto-oncogenes c-myc (*MYC*), *EGFR*, vascular endothelial growth factor c (*VEGFC*), Ras-like proto-oncogene A (*RALA*), casitas B-lineage lymphoma E3 ubiquitin ligase (*CBL*), and ETS proto-oncogene 1 (*ETS1*) (Fig. 3, Supplementary Figs. 9, 10). Importantly, activation of c-myc via the Wnt pathway can stimulate the cell cycle in part by activating expression or stimulating the accumulation of cyclin dependent kinase 6 (CDK6) and cell-cycle regular cyclin D2 (CCND2) in early G1 phase to promote transition to S phase to support proliferation^41–43^. *MYC*, *CDK6*, and *CCND2* transcripts were all significantly higher in R132Q HT1080* xenografts versus R132H (Supplementary Table 3). Though changes in *TP53* expression did not reach significance, tumor protein p53 inducible protein 3 (*TP53I3*) transcripts, which is induced by p53 and serves as a marker for pro-apoptosis^44^, were significantly lower in R132Q HT1080* xenografts than R132H (Supplementary Table 3). Indeed, transcripts of caspase-9 (*CAS9*), which drives apoptosis^45^, were marginally significantly lowered upon R132Q expression versus R132H with a *p* < 0.05 before correcting for multiple comparisons (Supplementary Table 3). Together, these data suggest that IDH1 R132Q-expressing chondrosarcoma models relied on upregulation of intra- and extra-cellular communication driven in part via direct CpG hypomethylation, as well as increased expression of oncogenes such as *MYC* and *EGFR* in mechanisms likely not directly dependent on altered DNA methylation.

### Epigenomic and transcriptomic features of mutant IDH1 glioma models

U87MG xenografts, which had more power than HT1080* xenografts due to higher tumor incidence, showed distinct clustering of IDH1 R132Q xenografts, with some clustering seen for WT and R132H tumors (Fig. 4a). PCA with overlaid D2HG levels showed better separation for the U87MG xenografts than HT1080* (Fig. 4b). Surprisingly, there was no observed increase in overall methylation among all CpG sites in IDH1 mutants compared to WT (Fig. 4c). Relative to the HT1080* xenografts, these mutant IDH1 U87MG xenografts had fewer differentially methylated CpG sites compared to WT (Fig. 4d). Therefore, we filtered the RRBS data and restricted our differential methylation analyses to variable CpG sites, and then found a large percentage of the differentially methylated sites were hypomethylated (n = 1,262) in R132Q-expressing U87MG tumor xenografts compared to both WT and R132H (Fig. 4f). Trends in location of hypermethylation and hypomethylation in the glioma models were similar to chondrosarcoma models, with most alterations occurring near promoter regions followed by distal intergenic and other intron regions (Fig. 4e). Compared to HT1080* xenografts, U87MG xenografts had more hypomethylation and hypermethylation in regions outside the CpG islands and shores in addition to the CpG islands (Supplementary Fig. 8).

**Fig. 4.**
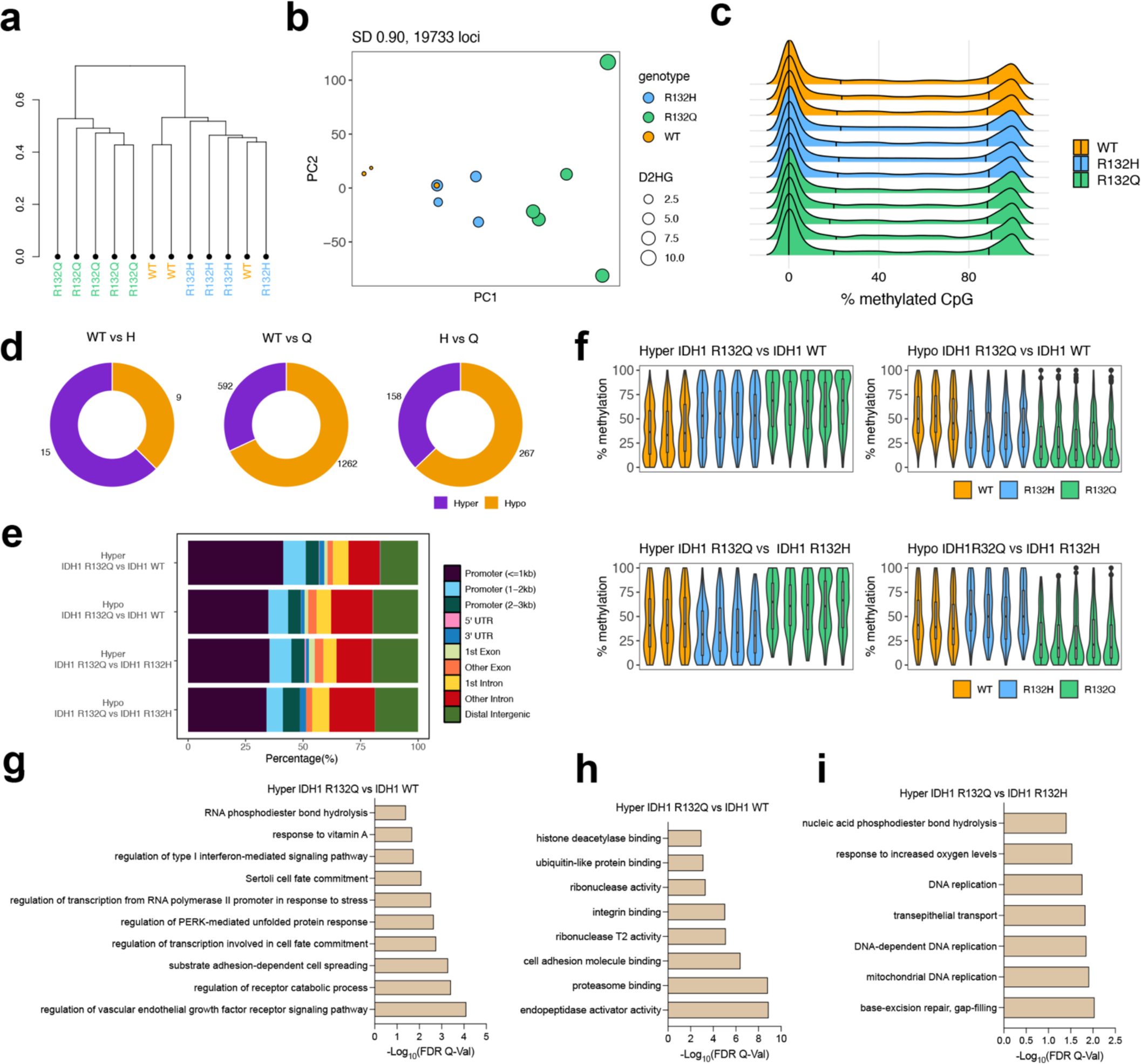
IDH1 mutants exhibit a subtler influence on the methylomes of U87MG tumors. Three of nine, four of nine, and five of nine (randomized selection for each) of the xenograft tumors that formed U87MG xenograft tumors upon IDH1 WT, R132H, or R132Q expression, respectively, were evaluated as biological replicates by RRBS. **a,** Hierarchical clustering of the most variable CpG sites (standard deviation > 0.90, n = 19,733) within the U87MG RRBS cohort. **b,** PCA plot of the most variable CpG sites (standard deviation > 0.90) within the U87MG RRBS cohort. The size of each dot corresponds to the level of D2HG (nmol tumor) measured in each tumor sample. Colors indicate genotype. **c,** Global methylation profiles of the U87MG RRBS tumor cohort showing percent methylation values of CpG sites for each sample. Black vertical lines indicate quantiles, colors indicate genotype. **d,** Number of differentially methylated CpG sites (DMS, % methylation difference ≥ 10 and q-value < 0.05) are shown. Of note, RRBS data is filtered to include variable loci (standard deviation > 10, n = 42,310) prior to differential methylation analysis. **e,** Distribution of the genomic features for the hyper- and hypo-methylated loci for all comparisons. **f,** Violin plots showing the distribution of percent (%) methylation of hypermethylated loci in IDH1 R132Q vs IDH1 WT (top left), hypomethylated loci in IDH1 R132Q vs IDH1 WT (top right), hypermethylated loci in IDH1 R132Q vs IDH1 R132Q (bottom left), and hypomethylated loci in IDH1 R132Q vs IDH1 R132H (bottom right) (n = 42,310, % methylation difference ≥ 10% and q-value < 0.05). For clarity, samples not included in the pairwise statistical comparisons are also shown. **g-i,** Gene ontology enrichment (using GREAT toolbox) showing enriched biological processes in hypermethylated (hyper) loci in IDH1 R132Q compared to IDH1 WT (**g**), molecular function in IDH1 R132Q vs IDH1 R132H (**h**), and biological processes in hypermethylated loci in IDH1 R132Q compared to IDH1 R132H (**i**).

### Pathways hypermethylated and down-regulated in IDH1 R132Q glioma models

RRBS analysis highlighted regions associated with nucleic acid phosphodiester bond hydrolysis and DNA replication and repair were hypermethylated in R132Q U87MG tumor xenografts versus R132H (Fig. 4i), reminiscent of DNA damage pathways hypermethylated in R132Q HT1080* tumors (Fig. 2k). This suggests that silencing DNA damage repair is an important strategy in both IDH1 R132Q-expressing tumor models. When comparing R132Q glioma xenografts with WT, pathways associated with cell adhesion, integrin binding, and stress response were highlighted as hypermethylated (Fig. 4g, 4h). Integrins, whose hypermethylation has been implicated in tumors^46^, mediate cell adhesion to ECM and their activation can induce intracellular signaling cascades to affect cell growth, migration, differentiation, adhesion, and apoptosis. RNAseq analysis showed no pathways being significantly downregulated upon R132Q expression versus R132H, though many similarities in downregulated pathways emerged for both mutants compared to WT, including pathways associated with chemotaxis, ECM organization, angiogenesis, and negative regulation of cell migration (Fig. 5, Supplementary Figs. 11, 12). Together, these data suggest that expression of both IDH1 mutants in glioma xenograft models share many transcriptionally down-regulated pathways, though IDH1 R132Q uniquely shows hypermethylation of DNA damage pathways.

**Fig. 5.**
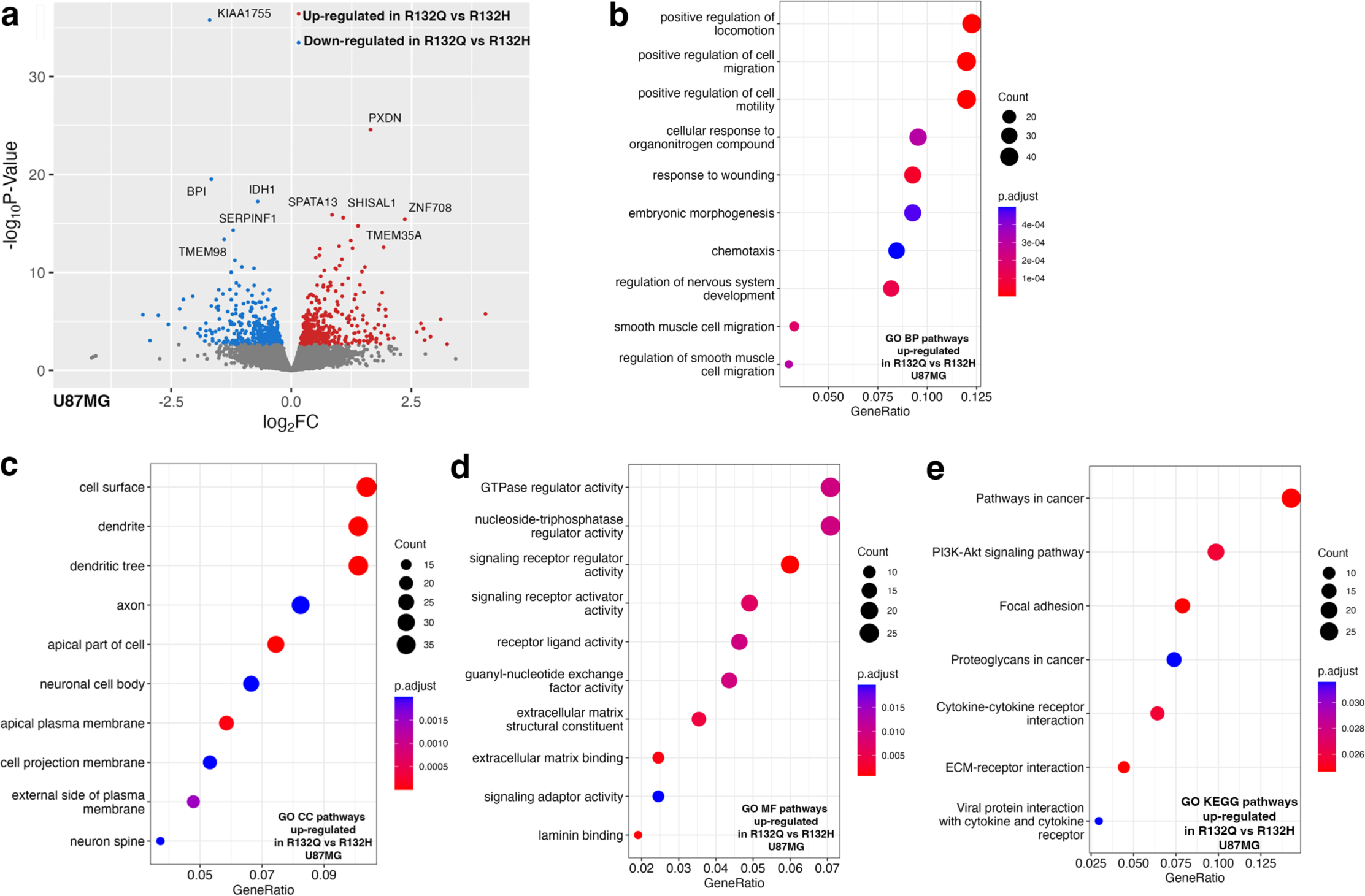
Transcriptome analysis using RNAseq of U87MG xenograft tumors comparing IDH1 R132Q and IDH1 R132H. Three of nine, four of nine, and five of nine (randomized selection for each) of the xenograft tumors that formed U87MG xenograft tumors upon IDH1 WT, R132H, or R132Q expression, respectively, were evaluated as biological replicates by RNAseq. Down-regulated pathways refer to categories where most genes show decreased expression levels in R132Q versus R132H IDH1. Up-regulated pathways refer to categories where most genes show increased expression levels in R132Q versus R132H IDH1. Number of genes is indicated in the count, with the *p*_adjusted_ value indicated by color. **a**, Volcano plot of differentially expressed transcripts comparing expression of IDH1 R132Q versus IDH1 R132H. **b**, Biological pathways (BP) up-regulated in IDH1 R132Q versus IDH1 R132H. **c**, Cellular component (CC) pathways up-regulated in IDH1 R132Q versus IDH1 R132H. **d**, Molecular function (MF) pathways up-regulated in IDH1 R132Q versus IDH1 R132H. **e**, KEGG pathways up-regulated in IDH1 R132Q versus IDH1 R132H.

### Pathways hypomethylated and up-regulated in IDH1 R132Q glioma models

RNAseq analysis showed that there were many upregulated pathways that may have helped drive the larger, earlier-forming tumors observed in IDH1 R132Q U87MG xenografts compared to R132H, including cell migration, motility, cell signaling pathways. A comparison of upregulated pathways in R132Q versus WT xenografts recapitulated reliance on these pathways, with transcripts of genes related to cell migration, motility, and negative regulation of differentiation upregulated (Supplementary Fig. 11). The PI3K/AKT/mTOR pathway emerged as a critical driver of these pro-tumorigenic features; PI3K/AKT/mTOR signaling via upregulation of *EGFR*, interleukin 6 (*IL6*), and janus kinase 1 (*JAK1*) were all significantly upregulated in R132Q U87MG xenografts versus R132H (Supplementary Table 4). Previously, the PI3K/AKT/mTOR pathway has been shown to be downregulated in R132H-driven tumors and tumor models. It was suggested that D2HG production inhibited AKT phosphorylation^47,48^, though upregulation in our R132Q xenografts suggests a different mechanism may be at play. *EGFR*, *IL6*, and *JAK1* upregulation also indicate JAK-STAT pathway activation upon R132Q expression, and we observed a nonsignificant trend of increased *STAT3* expression (Supplementary Table 4).

The PI3K/AKT/mTOR pathway, through activation of growth factor receptors like EGFR and integrins, can also activate MDM2, which in turn can degrade p53^49^. Notably, IDH1 R132Q-expressing U87MG tumor xenografts had significantly increased transcripts of many integrins, *EGFR*, and *MDM2* (Supplementary Table 4) compared to R132H. When comparing R132Q to WT, all transcripts except *MDM2* were also significantly increased (*MDM2* transcripts trended upwards), and *TP53* and *TP53I3* were significantly downregulated (Supplementary Table 4).

Notably, n-Ras (*NRAS*), as well as transforming growth factor β-1 (*TGFB1*) and transforming growth factor α (*TGFA*), which are downstream targets of n-Ras, were significantly upregulated in IDH1 R132Q-expressing U87MG xenografts compared to WT (Supplementary Table 4, Supplementary Fig. 11), indicating activation of cancer growth and metastasis pathways^50^. Many of these oncogenes are also implicated in focal adhesion, which helps regulate cell growth, migration, and ECM remodeling, and evidence of the importance of these pathways were supported by the upregulation of c-Jun (*JUN*), Rho associated coiled-coil containing protein kinase 2 (*ROCK2*), and several collagen genes upon R132Q expression compared to WT (Supplementary Table 4). Importantly, no significant upregulation in these oncogenes that R132Q appeared reliant upon was seen when comparing R132H versus WT (Supplementary Fig. 12, Supplementary Table 4). Together, these data highlight likely a non-methylation-based reliance of IDH1 R132Q-expressing U87MG xenografts on the activation of PI3K/AKT/mTOR, EGFR, and Ras pathways and deactivation of p53 to lead to increased cell growth and migration.

## Discussion

We expected that higher D2HG levels resulting from more efficient IDH1 R132Q catalysis would result in increased DNA methylation, but instead observed primarily genome hypomethylation in R132Q versus R132H tumors. Increased hypermethylation and hypomethylation are well-documented in R132H IDH1-mutated tumor models and tumor tissue^3,9,51–58^, though the mechanisms and consequences of hypomethylation are not understood^51–54^. Interestingly, malignant progression of mutant IDH1-driven lower grade gliomas was associated with partial DNA demethylation via an inability of the rapidly proliferating cells to maintain methylation^59^. Upon this passive demethylation, tumors had activated PI3K/AKT/mTOR and cell cycle pathways via mutation and altered transcript levels^59^, which is in contrast to the downregulation of MAPK, PI3K/AKT/mTOR, and EGFR pathways normally associated with IDH1 R132H-driven tumors^4^, with *EGFR* amplification and R132H IDH1 mutations are less common^60,61^ or even mutually exclusive^62^. Others have reported that faster progressing mutant-IDH1-driven astrocytomas had increased copy numbers in *EGFR*, *MYC*, *MET*, *CDK6*, and *CDKN2A/B* compared to slower-progressing patients^63^. Our observations of increased hypomethylation, larger tumor size, and activation of PI3K/AKT/mTOR, EGFR, Wnt, and Ras pathways upon R132Q expression supports these findings, suggesting that IDH1 R132Q confers features reminiscent of more aggressive, typically non-IDH1-driven tumors through mechanisms not likely directly involving CpG hypermethylation.

We found that correlation between DNA methylation and gene expression was relatively poor, and others have reported the same finding, citing that locations that acquire hypermethylation may often affect regions that already are transcriptionally less active^53^. Of course, changes in D2HG-driven histone methylation as well as indirect changes in protein activation via phosphorylation will also affect the transcriptome^4^. Our limited power in the WT IDH1 HT1080* tumors due to lack of tumor formation in 7/9 mice likely complicated analysis, as one of the WT tumors was unusually large (Supplementary Table 1).

The decreased methylation in mutant IDH1 U87MG xenografts compared to WT in our initial data processing was surprising. A possible reason for this is due to the low passage number in the U87MG cells; after successfully introducing the mutations, the U87MG cells were injected into mice at a passage number range of 6-12, while HT1080* lines were used at passage 21. We^9^ and others^64^ have shown that the degree of methylation changes over time, with high passage number associated with increased hypermethylation. However, deletion of IDH1 mutations have been reported both *in vitro* and *in vivo*^56^, and indeed, further expansion of our U87MG line, but not our HT1080*, led to the loss of expression of the R132Q mutation. To ensure that IDH1 R132Q was still expressed and to allow head-to-head comparisons of U87MG cells at similar passage number, xenograft injections occurred at this lower passage despite suboptimal conditions for RRBS analysis. Despite this challenge, a trend unique to IDH1 R132Q emerged in both chondrosarcoma and glioma xenografts – hypermethylation of pathways associated with DNA damage. There is support that hypermethylation of DNA damage pathways is unique to the IDH1 R132Q mutant; H3 histone hypermethylation studies with *in vivo* IDH1 R132H-expressing tumor models have suggested up-regulation of DNA damage response and cell cycle control^65^.

Altered Wnt signaling can ultimately result in dysregulation of differentiation and morphogenesis to help drive tumor progression^40,66–69^, with this pathway emerging as a possible therapeutic target due to its important role in glioma invasiveness and stemness^70^. Hypermethylation and/or down-regulation of Wnt signaling modulators and pathway members to result in aberrant Wnt signaling has been reported as a consequence of IDH1 R132H expression by us^9^ and others^53,71–73^. Hypermethylation of Wnt signaling in general has been implicated in a variety of cancers^67,74^, including hypermethylation of *WNT9A* and *FZD9* genes^66,75^. WNT9A has tumor suppressor features in that it drives the differentiation of chondrocytes^68^ and can inhibit cell proliferation^69^. We found here that alterations in the Wnt pathway represented another persistent phenotype in our xenograft models, with primarily hypermethylation (and some hypomethylation) of gene pathway members upon expression of IDH1 R132Q in chondrosarcoma models. Notably, transcripts of many Wnt pathway members were significantly up- and down-regulated upon IDH1 R132Q expression in both xenograft models (Supplementary Tables 3, 4), suggesting that non-epigenetic changes may also alter the regulation of this pathway.

Finding unique methylation and transcription patterns when comparing HT1080* and U87MG xenografts was unsurprising, as this has been reported in tumors and tumor models of mutant IDH1-driven cancers. HT1080 and U87MG lines represent different cells of origin, and HT1080* cells used here originated from an IDH1^R132C/+^ background that likely had epigenetic changes prior to deletion of the R132C allele. HT1080* cells also have only half the endogenous WT allele amount compared to U87MG cells, and there have been differences reported between hemizygous and heterozygous tumor models^76^. Interestingly, there is a clear mutation selection bias in gliomas (IDH1 R132H) versus chondrosarcomas (IDH1 R132C)^12^. Though differences in catalytic profiles between these two mutants were not nearly as profound as that seen in R132Q, we showed previously that R132C has a ∼3-fold increase in *k*_cat_ compared to R132H^20^. Thus, this preference for the R132C mutation in chondrosarcomas versus R132H in gliomas raises the possibility that glioma cells could be more susceptible to D2HG toxicity and thus prefer the lower D2HG levels associated with R132H. There is debate about which mutant IDH-driven tumor type is associated with more hypermethylation. Some have shown that IDH-driven gliomas are more hypermethylated than AML, melanomas, and cholangiocarcinomas^54^, though others report AML as having the highest methylation^51^. Understanding the unique DNA and histone methylation features among different tumor tissue and among different IDH1 and IDH2 mutants remain important areas of research.

In summary, we leveraged steady-state kinetics, metabolic profiling, methylomics, and transcriptomics to connect the biophysical features of tumor-driving IDH1 mutations to *in vivo* phenotypes in tumor models. Identification of an IDH1 mutation with distinctly robust neomorphic activity provided an ideal opportunity to establish the consequences of IDH1 mutation with low versus high catalytic efficiency for D2HG production. Our findings showed that higher catalytic activity afforded by R132Q versus R132H led to increased D2HG levels in cells, tumor tissue, and sera, and larger tumors upon expression of this highly kinetically active mutant, despite its lower expression. Further, IDH1 R132Q expression led to activation of several pro-tumor pathways, including those associated with EGFR and PI3K signaling reminiscent of lower grade mutant IDH1 tumors that progress to higher grade. Our work highlights a potential ability of D2HG levels to tune phenotype severity through mechanisms beyond increased DNA hypermethylation.

## Online Methods

### Chemicals and reagents

β-Nicotinamide adenine dinucleotide phosphate reduced trisodium salt (NADPH), β-nicotinamide adenine dinucleotide phosphate disodium salt (NADP^+^) tris(2-carboxyethyl)phosphine) (TCEP), agarose, *L*-norvaline, and 100 kD Amicon centrifugal filters were purchased from Millipore Sigma (Burlington, MA). Triton X-100, magnesium chloride (MgCl_2_), α-ketoglutaric acid sodium salt (αKG), DL-isocitric acid trisodium salt hydrate, dithiothreitol (DTT), isopropyl 1-thio-β-D-galactopyranoside (IPTG), Pierce protease inhibitor tablets, chloroform, isopropanol, PEI STAR transfection reagent, chloroquine diphosphate, T-PER Tissue Protein Extraction Reagent, β-tubulin polyclonal antibody, Pierce ECL Western blotting substrate, and H3K9me3 polyclonal antibody were purchased from Thermo Fisher Scientific (Waltham, MA). Β-mercaptoethanol (BME) was bought from MP Biomedicals (Santa Ana, CA). Nickel-nitrilotriacetic acid (Ni-NTA) resin was obtained from Qiagen (Valencia, CA), and stain-free gels (4-12%) were purchased from Bio-Rad Laboratories (Hercules, CA). High glucose Dulbecco’s Modified Eagle’s Medium (DMEM), phosphate-buffered saline (PBS), Penicillin-streptomycin-glutamine (PSG) 100X, L-glutamine, and TrypLE express enzyme were purchased from Gibco (Grand Island, NY). Fetal bovine serum (FBS) was obtained from Cytiva (Marlborough, MA). Matrigel was purchased from Corning (Corning, NY). Recombinant anti-HA-tag and anti-γ-H2A.X antibodies were purchased from Abcam (Waltham, MA). Pre-cast 4-20% gels, polyvinylidene fluoride (PVDF) membranes, Precision Plus Protein Dual Color Standards ladder, and 4X Laemmli sample buffer were purchased from Bio-Rad Laboratories (Hercules, CA). MEGAPREP 3 kit was purchased from Bioland Scientific (Paramount, CA). Goat anti-rabbit IgG (H+L) HRP was purchased from Invitrogen (Carlsbad, CA). Anti-mouse IgG HRP-linked antibody was purchased from Cell Signaling Technology (Danvers, MA). GentleMACS M Tubes were purchased from Miltenyi Biotec (San Diego, CA).

### Protein expression, and purification

Human IDH1 WT, R132H, and R132Q cDNA constructs in a pET-28b(+) plasmid were transformed in *E. coli* BL21 Gold DE3 cells. After incubation in 0.5-1 liters of terrific broth containing 30 µg/mL of kanamycin (37°C, 200 rpm), induction began with 1 mM IPTG (final concentration) after briefly cooling cultures to 25 °C after reaching an A_600_ of 0.9-1.2. After 18 h incubation (19 °C, 130 rpm), cell pellets were harvested and resuspended in lysis buffer (20 mM Tris pH 7.5 at 4°C, 500 mM NaCl, 0.1% NaCl, 0.1% Triton X-100, and 1 protease inhibitor tablet) for cell lysis via sonication. Crude lysates were clarified via centrifugation at 12,000 rpm for 1h. The resulting lysate was loaded on to a pre-equilibrated Ni-NTA column, followed by 150 mL of wash buffer (20 mM Tris pH 7.5 at 4 °C, 500 mM NaCl, 15 mM imidazole, 5 mM BME). Protein was then eluted with elution buffer (50 mM Tris pH 7.5 at 4°C, 500 mM NaCl, 500 mM imidazole, 5% glycerol, 10 mM BME). Protein was concentrated if needed and dialyzed overnight in 50 mM Tris pH 7.5 @ 4°C, 100 mM NaCl, 20% glycerol, and 1 mM DTT. > 95% purity was ensured via SDS-PAGE analysis, and IDH1 was flash frozen using liquid nitrogen, and stored at -80 °C. All kinetic analysis was performed < 1 month from cell pelleting.

### Enzyme kinetic analysis

An Agilent Cary UV/Vis 3500 spectrophotometer (Santa Clara, CA) was used to perform steady-state kinetic assays at 37 °C. To allow heterodimers to form, a 1:1 mixture (200 nM final concentration of both WT and mutant IDH1) was incubated on ice for 1 h after gentle mixing. However, we note that a mixture of homodimers and heterodimers, depending on their binding equilibria, may exist. To measure the conventional reaction (conversion of ICT to αKG), a cuvette containing IDH1 assay buffer (50 mM Tris, pH 7.5 at 37 °C, 150 mM NaCl, 10 mM MgCl_2_, 1 mM DTT), and the mixture of WT and mutant (R132H or R132Q) IDH1 (200 nM) were preincubated for 3 min at 37 °C. NADP^+^ (200 µM) and varying concentrations of ICT were added to initiate the reactions and the change in absorbance due to NADPH formation was monitored at 340 nm. The same conditions were used for the neomorphic reaction (conversion of αKG to D2HG) except NADPH and αKG were used to initiate the reaction and the consumption of NADPH was monitored. The pH of αKG was adjusted to 7.0 prior to use. The slope of the linear range of the change in absorbance over time was calculated and converted to nanomolar NADPH using the molar extinction coefficient for NADPH of 6.22 cm^-1^ mM^-1^ to determine *k*_obs_ (i.e. nM NADPH/nM enzyme s^-1^) at each substrate concentration. Each *k*_obs_ was fit to the Michaelis-Menten equation in GraphPad Prism (GraphPad Software, La Jolla, CA) to calculate *k*_cat_ and *K*_m_. In all cases, two biological replicates (2 protein preparations) were used to measure *k*_obs_ over multiple days, ensuring both preparations were tested at the full range of concentrations of substrate to ensure batch-to-batch reproducibility. Kinetic parameters are reported as standard error (S.E.) resulting from deviation from the mathematical fit of the equation.

### Cell line generation and culturing

LEIH-IDH1-R132H, LEIH-IDH1-WT, and LEIH-IDH1-EV plasmids conferring hygromycin resistance and containing a *C*-terminal HA-tag were obtained from William Kaelin Jr. (Dana Farber Cancer Institute). The LEIH vector has an EF1a promoter for high expression. We used site-directed mutagenesis to introduce the R132Q mutation by removing the IDH1 WT insert from the LEIH vector, inserting this into a pET17b vector and performing mutagenesis using a KAPA HiFi Hot Start Ready Mix PCR kit (Roch, Boston, MA) using the following mutagenesis primers: forward -- 5’-CTGTATTGATCCCCATAAGCATGCTGACCTATGATGATAGGTTTTACC; reverse – 5’-GGTAAAACCTATCATCATAGGTCAGCATGCTTATGGGGATCAATACAG. The IDH1 insert was then returned to the LEIH vector. All LEIH plasmids confer kanamycin resistance in bacteria and were amplified in DH5α (Thermo Scientific, Fisher Waltham, MA). Plasmids were purified using MEGAPREP 3 (Bioland Scientific, Paramount, CA). Plasmid whole genome sequencing was performed by Retrogen Inc. (San Diego, CA). The hygromycin-resistant genes linked to IDH1 via the internal ribosome entry site (IRES) allow its continuous growth in a low concentration of hygromycin, which was supplemented to the media at the selection dose of 100 µg/mL (final concentration). Only cells that survived were selected, and fresh media was added to flasks with a concentration of 60 µg/mL hygromycin to prevent silencing of the mutation over time. Lentiviral particles were produced in HEK293T Lenti-X cells (Clontech, Mountain View, CA). Cells (5×10^6^) were seeded in a 15 cm culture flask and allowed to reach ∼70% confluency. LEIH vectors of interest were co-transfected with the lentiviral packaging, envelope, and infection system plasmids pVSVG, pRev, and pGal/Pol (Addgene, Watertown, MA). The transfected cells were incubated for 16 h, and the media was collected and passed through a 45 µM vacuum filter (Corning, Corning, NY). To concentrate the viral particles, centrifugation of the sample in a 100kD Amicon centrifugal filter was used (Sigma Aldrich, St. Louis, MO), and the viral particles were stored at -80 °C.

U87MG cells were purchased from ATCC (Manassas, VA). HT1080 cells, which contain an endogenous R132C mutation (IDH1^R132C/+^) were modified by Kun-Liang Guan (University of California, San Diego) to knock-out the mutant allele to generate (IDH1^-/+^)^77^, here notated as HT1080*, which were then provided to us. Normal human astrocytes (NHA) were obtained from Russ Pieper (University of California, San Francisco), and C28 cells were obtained from Johnathan Trent (University of Miami). Routine mycoplasma testing (PCR mycoplasma detection kit, Thermo Scientific, Waltham, MA) was performed as infrequently as every 6 months and as frequently as monthly. Cells stably over-expressing IDH1 WT, R132H, or R132Q were prepared in each of these four cell lines by culturing cells in T75 flasks and allowing them to reach ∼60% confluency in DMEM (Gibco, Grand Island, NY) supplemented with 10% FBS. Subsequently, the media was replaced with fresh DMEM, and 20 µL of a fresh aliquot of LEIH virus was added directly to the flasks. After 72 h past transfection, media was replaced with DMEM containing 10% FBS and 60 µg/mL hygromycin for cell selection. To assess the success of transfection, we extracted the total protein from the mutant cells and conducted western immunoblotting using a recombinant anti-HA tag antibody (Abcam, Waltham, MA).

For routine culturing, the U87MG, HT1080*, NHA, and C28 cells were cultured in DMEM (Gibco, Grand Island, NY) containing 10% FBS (Cytiva, Marlborough, MA) and hygromycin at 60 µg/mL (final concentration). HT1080 and U87MG cells were seeded in the concentration of 2×10^5^ of U87MG in a T75 flask. Cells were allowed to reach ∼60% confluency before splitting, with fresh media was added every 48 h.

### Ethics Statement

All animal experiments were conducted in accordance with the San Diego State University (SDSU) Institutional Animal Care and Use Committee (protocol approval number 22-06-007s).

### Animals

Female athymic nude mice (6-8 weeks old) were purchased from Jackson Labs (Sacramento, CA). Female mice were selected based on previously reported protocols for mutant IDH1 xenograft generation^78^. Mice underwent a mandatory period of 1-week acclimatization at the SDSU Animal Facility. The animals were housed in individually ventilated cages (ABSL2) with a maximum of 4 mice per cage. The mice housing environment was maintained throughout the study with a temperature range of 22 to 25°C, humidity between 40 to 60%, and a light cycle of 12 hours of light and 12 hours of darkness. Cages, bedding, water, and food were individually irradiated for sterility. Animal procedures and maintenance were carried out in accordance with SDSU Institutional Animal Care and Use Committee guidelines.

### Xenograft generation and analysis

For xenografts, 2×10^6^ U87MG IDH1 WT, IDH1 R132H, or IDH1 R132Q cells suspended in Matrigel (354263 Corning, Corning, NY) and PBS (Gibco, Grand Island, NY) (1:1) or 2 × 10^6^ HT1080* IDH1 WT, IDH1 R132H, or IDH1 R132Q cells suspended in PBS were implanted subcutaneously into the left flanks of 6–8-week-old female athymic Nu/Nu mice (Jackson Laboratory). For each IDH1 expression condition (WT, R132H, or R132Q), 9 mice were used (27 mice total). Based on previous work on generating xenografts with HT1080 cells^79,80^, Matrigel was not used for the HT1080* cells. Vehicle controls (1:1 Matrigel/PBS or PBS alone) were injected subcutaneously into the corresponding right flanks. Twice weekly, mice were weighed, and tumor volume was measured with a Vernier caliper and calculated using the following equation: Tumor Volume (mm^3^) = ½ length^2^ (long axis) × width (short axis). Mice were sacrificed once tumors reached 20 mm in length or 120 days of growth. Blood was collected in BD microtainer SST tubes (BD Franklin Lakes, NJ) and processed according to manufacturer’s protocol. Tumors were weighed. Serum and tumor tissue were flash-frozen for downstream analysis.

### Western immunoblotting

Antibodies EGFR (catalog # 4267S, lot 24), p-EGFR (catalog # 3777T, lots 17 and TBD), JAK1 (catalog # 3332S, lot 6), STAT3 (catalog # 9139S, lot TBD), p-STAT3 (catalog # 9145S, lot TBD), ATM (catalog # 2873S, lot 7), p-ATM (catalog # 13050S, lot 6), Chk2 (catalog # 2662S, lot 9), p-Chk2 (catalog # z82263S, lot 13), Rb (catalog # 9309S, lot 14), p-Rb (catalog # 8180S, lot 7), were purchased from Cell Signaling Technology (Danvers, MA). Antibodies HA (catalog #PA1-985 lots XG358301 and YL378360), HRP-rabbit (catalog # 3179S, lot 21), and HRP-mouse (catalog # 3179S, lot 21) were purchased from Invitrogen (Waltham, MA). GAPDH antibody (catalog # MAB374, lot 3988901) was purchased from Sigma-Aldrich (St. Louis, MO).

Protein was extracted from tumor tissue by combining 100–200 mg of tumor tissue with 500 µL of T-PER in gentleMACS M tubes, and samples were dissociated mechanically using a gentle MACS Dissociator (Miltenyi Biotec, San Diego, CA). All tubes were incubated on ice for 15 min post-tissue homogenization. After incubation, samples were transferred to autoclaved 1.7 mL microtubes and clarified by centrifugation at 13,000 rpm for 20 min. The supernatant was then transferred to fresh 1.7 mL microtubes and stored at -20 °C. Protein was extracted from cell lines by culturing cells in T75 flasks and allowed them to reach ∼60% confluency in DMEM + 10% FBS. Cells were washed twice with PBS and detached from the flask using TrypLE Express enzyme. The enzyme activity was neutralized using complete media (1:1). Cells were transferred to 15 mL conical and clarified using centrifugation at 1000 rpm for 10 min, after which the supernatant was discarded. Homemade RIPA buffer (200 µL of 10 mM Tris pH 8.0, 140 nM NaCl, 1 mM EDTA, 0.5 mM EGTA, 1% Triton X-100, 0.1% sodium deoxycholate, and 0.1% SDS) treated with a Pierce Protease Inhibitor Mini Tablet was added to the cell pellet supplemented and incubated on ice for 15 min. The cell lysates were then transferred to an autoclaved 1.7 mL microtube and clarified by centrifugation at 3,000 rpm for 20 min. All cell line protein samples were stored at -20 °C. In all cases, protein samples were normalized to 20 µg/mL of protein in 1X NuPAGE LDS sample buffer and Bond-Breaker TCEP solution, and incubated at 95 °C for 10 min. Protein samples were loaded to a NuPAGE 4–12% bis-tris precast gel for sample separation at 130 V for 90 min following a pre-electrophoresis at 20 V for 10 min. Protein samples were transferred to PVDF Immobilon-FL, 0.45 µm membrane using an XCell SureLock Blot Module for 60 min at 1.0 A, 30 V. Following blocking membranes in 5% skim milk in 0.1% PBST, or 5% BSA in 0.1% PBST the following antibody concentrations were used: anti-HA-tag (1:1000), γ-H2A.X (1:1000), GAPDH (1:6000), and β-tubulin (1:5000). Following washing, secondary antibodies were added at the following concentrations: goat anti-rabbit IgG (H+L) HRP (1:2000) and anti-mouse IgG, HRP-linked (1:2000). Following washing in PBST, membranes were treated with SuperSignal West Pico PLUS Chemiluminescent Substrate for imaging on an iBright CL1000 Imaging Systems (ThermoFisher).

### Metabolite analysis by mass spectrometry

Three biological replicates of U87MG, HT1080*, C28, and NHA cells were allowed to reach ∼60% confluency in DMEM supplemented with 10% FBS. After washing the cells three times with cold PBS, 0.45 mL of 20 µM *L*-norvaline in 50% methanol (50% v/v in water) was added to serve as an internal standard. Flasks were incubated on dry ice for 30 min and then thawed on ice for 10 min. Cells were lifted from the flask using a cell scraper and transferred to an eppendorf tube. For mouse studies, for each biological replicate, ∼20 mg of each tumor xenograft was cut and weighed, and serum was collected. All samples were flash frozen in polypropylene tubes. All cell, tumor, and serum samples were shipped in dry ice to Sanford Burnham Prebys Medical Discovery Institute Protein Production and Analysis Facility to perform metabolite measurements. Dried methanol extracts were first derivatized by adding 50 μl 20 mg/mL methoxyamine-hydrochloride prepared in dry pyridine (Sigma Aldrich St. Louis and Burlington, MA) and incubation for 20 min at 80 °C. After cooling, 50 μL *N*-tert-butyldimethylsilyl-*N-*methyltrifluoroacetamide (Sigma Aldrich) was added, and samples were re-incubated for 60 min at 80 °C before centrifugation for 5 min at 14,000 rpm at 4 °C. The supernatant was transferred to an autosampler vial for gas chromatography-mass spectrometry (GC-MS) analysis. A TSQ 9610 GC-MS/MS (Thermo Scientific) with an injection temperature of 250 °C, injection split ratio 1/10, and injection volume 0.5 μL was used. GC oven temperature started at 130 °C for 4 min, rising to 243 °C at 6 °C/min and to 280 °C at 60 °C/min with a final hold at this temperature for 2 min. The GC flow rate with helium carrier gas was 50 cm/s, and a GC 15 m × 0.25 mm × 0.25 μm SHRXI-5ms column was used (Shimadzu, Columbia, MA). GC-MS interface temperature was 300 °C, and the ion source temperature was 200 °C, with 70 V/150 μA ionization voltage/current. The mass spectrometer was set to scan m/z range 50–600, with ∼1 kV detector sensitivity. For analysis of pyruvate, the derivatization procedure was modified by the substitution of ethylhydroxylamine for methoxyamine, and the initial GC-MS oven temperature was 110 °C for 4 min, rising to 230 °C at 6 °C/min and after that as above. Metabolites were quantified against varied amounts of standard mixtures run in parallel and data were analyzed using Metaquant v1.3.5. Quantities were corrected for recovery using the *L*-norvaline internal standard. These methods have been previously reported^81^.

### PCA analysis

Dimensionality reduction method was carried out on all samples using Principal Component Analysis (PCA) in RStudio 2023.06.0 Build 421. Metabolites were read as .csv files and assigned as data matrices and PCA was carried out on sample groups using the *prcomp* function. Missing values were removed using *na.omit* and PCA plots were generated using ggplot2^82^.

### RRBS

100–200 mg of tumor tissue from 12 U87MG tumor xenografts (3 WT, 4 R132H, and 5 R132Q tumors, all randomly selected) and 9 HT1080* tumor xenografts (2 WT, 3 R132H with one failed data collection that was then eliminated, and 5 R132Q tumors, all randomly selected) was weighed and then shipped in dry ice for RRBS analysis at Active Motif (Carlsbad, CA). There, tissues were subjected to proteinase K digest solution (0.5% SDS, 0.5mg/ml proteinase K, 100 mM EDTA, in TE pH 8) by allowing the samples to rotate overnight at 55⁰C. For library preparation and sequencing, 100 ng of gDNA was digested with TaqaI (NEB Ipswich, MA) at 65 ⁰C for 2 h followed by MspI treatment (NEB Ipswich, MA) at 37⁰C overnight. Following enzymatic digestion, samples were used for library generation using the Ovation RRBS Methyl-Seq System (Tecan, Männedorf, Switzerland) following the manufacturer’s instructions. The digested DNA was randomly ligated, and, following fragment end repair, bisulfite was converted using the EpiTect Fast DNA Bisulfite Kit (Qiagen 59824) following the Qiagen protocol. After conversion and clean-up, samples were amplified, resuming the Ovation RRBS Methyl-Seq System protocol for library amplification and purification. Libraries were measured using the Agilent 2200 TapeStation System and quantified using the KAPA Library Quant Kit ABI Prism qPCR Mix (Roch, Boston, MA). Libraries were sequenced on a NovaSeq 6000 (Illumina, San Diego, CA) at SE75 x 30M reads (single-end).

### RRBS data processing and analysis

Methylation calls were produced using Bismark and imported into the MethylKit R package (v. 1.28.0)^83^. CpGs with more than ten reads were included in the downstream analysis. Filtered counts were normalized and CpG dinucleotides were merged (destrand = FALSE). Most variable CpGs across tumors were calculated as having a standard deviation > 0.90. Samples were visualized using PCA and hierarchical clustering using the Ward method. For HT1080* tumors, to perform pairwise comparisons in methylation levels, samples of interest were merged into one object containing common CpGs. Pearson correlation was used to assess concordance between D2HG amount in the tumors and DNA methylation levels of CpG sites. Differential methylation analysis was performed using Fisher’s exact test and differentially methylated CpGs were selected based on q < 0.05 and percent methylation difference > 20%. For U87MG tumors, CpGs were first filtered to include the most variable sites (CpGs with standard deviation > 10) and differential methylation analysis was performed as described, with a notable exception that CpGs were selected based on q < 0.05 and methylation difference > 10%. Feature annotation was done using ChIPseeker (v. 1.38.0) R package^84,85^. Genomic annotations for hg38 were retrieved using the TxDb.Hsapiens.UCSC.hg38.knownGene R package^86^. GREAT analysis software tool (with default parameters)^87,88^ was applied to determine enriched GO terms near the differentially methylated CpGs. The entire filtered CpG sites used for differential methylation analysis was used as the genomic background for GREAT analysis.

### RNA-seq

RNA samples were processed at the University of California, San Diego Institute for Genomic Medicine (IGM). RNA samples were extracted from 12 U87MG tumor xenografts (3 WT, 4 R132H, and 5 R132Q tumors, all randomly selected and identical to those selected for RRBS analysis) and 10 HT1080* tumor xenografts (2 WT, 3 R132H, and 5 R132Q tumors, all randomly selected and identical to those selected for RRBS analysis) using QIAzol Lysis Reagent (QIAGEN, Germantown, MD). 100–200 mg of the tumor tissue was placed in gentle MACS M tubes (Miltenyi Biotec San Diego, CA). 500 µL of QIAzol Lysis Reagent was added to the tissue, and the samples were dissociated mechanically using a gentle MACS Dissociator (Miltenyi Biotec, San Diego, CA). All tubes were incubated at room temperature for 5 min, and cell lysate was transferred to an autoclaved Eppendorf tube. Chloroform (0.2 mL) was added, followed by 30 s vortexing and a 5 min incubation at room temperature. Following separation by centrifugation at 12,000 rpm for 15 min at 4 °C, the top of three total layers containing RNA was transferred to a new autoclaved Eppendorf tube. After adding 0.5 mL of isopropanol samples were mixed by vortexing for 15 s and incubated at room temperature for 10 min. Tubes were then clarified by centrifugation at 12,000 rpm for 10 min at 4 °C. The supernatant was aspirated and 1 mL of 75% ethanol was added to the pellet that was mixed with vortexing for 15-30 s. Samples were then separated by centrifugation at 7,500 rpm for 5 min at 4 °C. The supernatant was removed, and the RNA pellets were air-dried for 5 min and then dissolved in 50-70 µL of RNase-free water. The integrity of RNA was assessed on a 1% agarose gel in 1x TAE buffer (0.4 M tris acetate pH 8.3 at room temperature, 0.01 M EDTA). An RNA quality control check was conducted at IGM and only high-quality RNA samples with RIN ≥ 7.4 were used to generate the RNA-seq libraries. Libraries were generated using the Illumina Ribo-Zero Plus rRNA Depletion Kit with IDT for Illumina RNA UD Indexes (Illumina, San Diego, CA). All samples were processed according to the manufacturer’s instructions. Generated libraries were multiplexed and sequenced with 150 base pair (bp) Paired End (PE150) to a depth of approximately 25 million reads per sample on an Illumina NovaSeq 6000. Samples were finally demultiplexed using bcl2fastq v2.20 Conversion Software (Illumina, San Diego, CA).

### RNA-seq data processing and analysis

The quality of the raw RNAseq data was evaluated using FASTQC v0.12.1^89^. Based on the FASTQC reports, potential adapter contamination and low-quality sequences were identified and trimmed using Cutdapt v4.8^90^. Cutadapt parameters were set for paired-end reads, with a quality cutoff of 20 and a minimum filter length of 20. The genome index was constructed using the hg38 reference genome assembly obtained from the National Center for Biotechnology Information (NCBI) with the Spliced Transcripts Alignment to a Reference (STAR) aligner version v2.7.11b^91^. Subsequently, quality-filtered reads were aligned to this genome index using STAR. The aligned reads were sorted by read name using sambamba v1.01 to prepare them for counting^92^. featureCounts v2.0.6 was utilized to count the number of reads per annotated gene^93^. The raw gene counts were normalized using DESeq2 to ensure differences in sample read counts represented biological differences rather than technical variation^94^. DESeq2 was also used to identify differentially expressed genes (DEGs) in the mutants when compared to the WT form. Significant DEGs were defined as genes exhibiting an adjusted p-value (FDR) below 0.05. Pathway analysis was performed to identify significantly enriched gene sets from the Kyoto Encyclopedia of Genes and Genomes (KEGG), as well as the Gene Ontology (GO) databases covering biological processes, molecular functions, and cellular components using ClusterProfiler v4.10.1^95–97^. Significantly enriched, upregulated, and downregulated pathway alterations between sample groups were determined by an adjusted *p*-value (FDR) below 0.05.

## Data availability

The complete set of RNAseq and RRBS data can be acquired via NCBI’s Gene Expression Omnibus (GEO) under accession numbers upon publication. Supplementary Information is included with Supplementary Figs and Tables. Additional information and requests for resources and reagents should be directed for fulfillment to corresponding author Christal D. Sohl.

## Acknowledgments

LEIH vectors containing WT and R132H IDH1 were a gift from William Kaelin Jr. (Dana Farber Cancer Institute). HT1080* were a gift from Kun-Liang Guan (UC San Diego). Normal human astrocytes (NHA) were a gift from Russ Pieper (UC San Francisco). C28 cells were a gift from Johnathan Trent (University of Miami). This work was funded by a Research Scholar Grant, RSG-19-075-01-TBE, from the American Cancer Society (C.D.S.), National Institutes of Health R35 GM137773 (C.D.S.) and R25 GM058906 (SDSU), the California Metabolic Research Foundation (SDSU), the German Cancer Aid Max Eder Program grant, 70114934 (S.T.), and the Rees-Stealy Research Foundation (E.A.). This publication includes data generated at the Sanford Burnham Prebys Protein Production and Analysis Facility, which is supported by NCI Cancer Center Support Grant P30 CA030199, and from the UC San Diego IGM Genomics Center, who utilized an Illumina X Plus that was purchased with funding from a National Institutes of Health SIG grant (#S10 OD026929). The RRBS data collection reported here was obtained at Active Motif (Carlsbad, CA).

## Author contributions

M.A.A.A. performed culturing, sample preparations, and aided in xenograft characterization. M.R. performed the xenograft work, data analysis, and created visualizations. A.S. performed RNAseq analysis and created visualizations. G.W. optimized culturing conditions and performed data analysis. A.H. performed culturing and aided in cell line generation. E.A. performed the protein expression and purification and kinetic analysis. C.G. aided in xenograft characterization, data analysis, and created visualizations. J.W. aided in xenograft characterization. G.Q. aided in protein expression. U.Z.G., C.D.H., S.T., and C.D.S. contributed to study supervision. S.T. performed RRBS analysis, created visualizations, and helped write the manuscript. C.D.S. designed the study, performed data analysis, created visualizations, and wrote the manuscript. All living authors contributed to editing.

## Additional information

Supplementary information is included.

Correspondence and requests for materials should be addressed to C.D.S.

