## Supplementary Information for "Catalytically distinct IDH1 mutants tune phenotype severity in tumor models"

Supplementary Figs. 1-12

Supplementary Tables 1-4

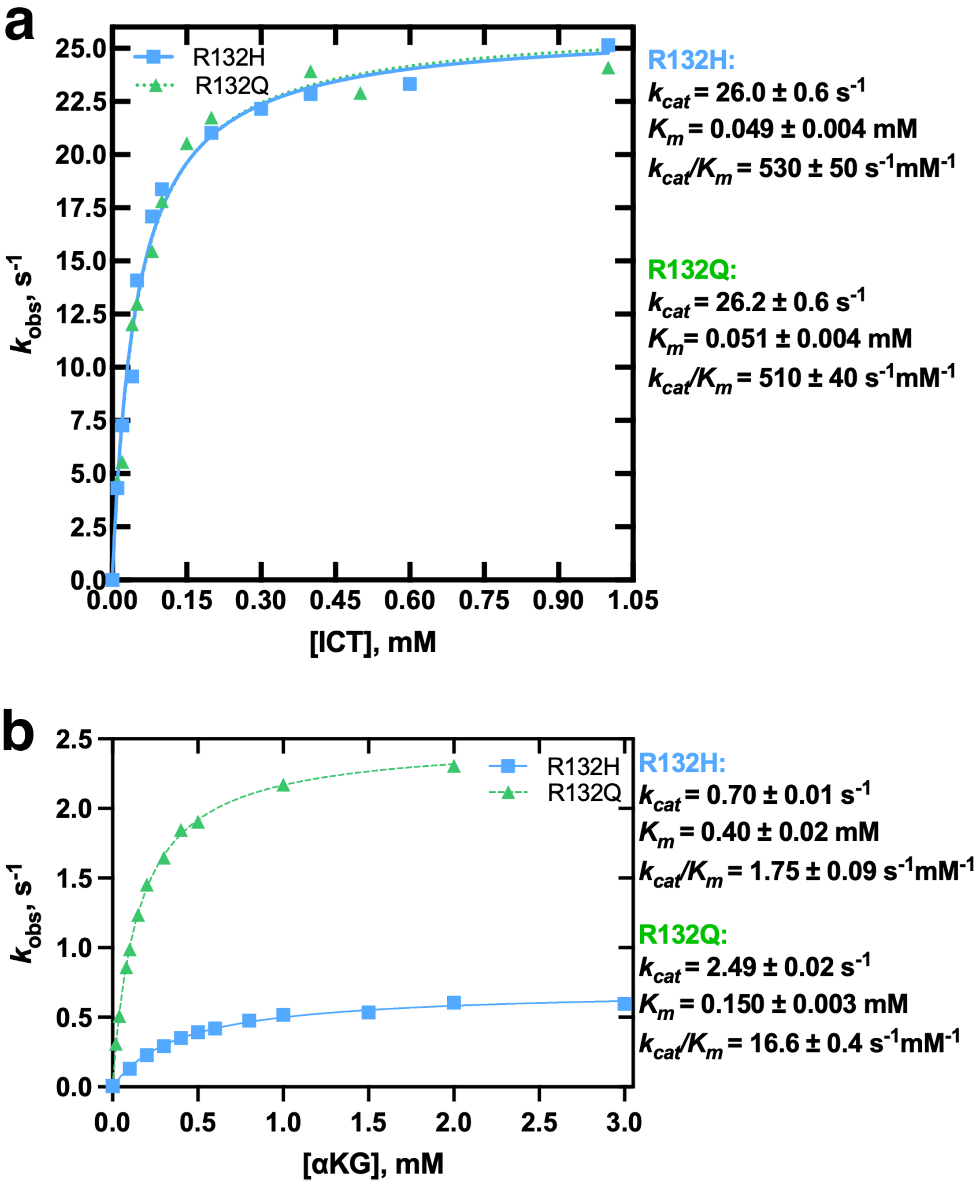


**Supplementary Fig. 1. Steady-state kinetic analysis for WT and R132H and R132Q mutant mixtures.** Shown is a 1:1 mixture of WT and mutant IDH1 to allow heterodimerization to occur, though populations of WT:WT and mutant:mutant homodimers may also exist. **a**, Steady-state kinetic parameters for the conventional reaction of the conversion of ICT to αKG was measured as a function of varying substrate concentration The following kinetic parameters were measured for a mixture of WT and R132H: $k_{cat,\ ICT\rightarrow\alpha KG}$ = 26.0 ± 0.6 $s^{-1}$; $K_{m,\ ICT}$ = 0.049 ± 0.004 mM; $k_{cat}/K_{m,\ ICT\rightarrow\alpha KG}$ = 530 ± 50 $mM^{-1}$ $s^{-1}$. The following kinetic parameters were measured for a mixture of WT and R132Q: $k_{cat,\ ICT\rightarrow\alpha KG}$ = 26.2 ± 0.6 $s^{-1}$; $K_{m,\ ICT}$ = 0.051 ± 0.004 mM; $k_{cat}/K_{m,\ ICT\rightarrow\alpha KG}$ = 510 ± 40 $mM^{-1}$ $s^{-1}$. **b**, Steady-state kinetic parameters for the neomorphic reaction of the conversion of αKG to D2HG were measured as a function of varying substrate concentration. The following kinetic parameters were measured for a mixture of WT and R132H: $k_{cat,\ \alpha KG\rightarrow D2HG}$ = 0.70 ± 0.01 $s^{-1}$; $K_{m,\ \alpha KG}$ = 0.40 ± 0.02 mM; $k_{cat}/K_{m,\ \alpha KG\rightarrow D2HG}$ = 1.75 ± 0.09 $mM^{-1}$ $s^{-1}$. The following kinetic parameters were measured for a mixture of WT and R132Q: $k_{cat,\ \alpha KG\rightarrow D2HG}$ = 2.49 ± 0.02 $s^{-1}$; $K_{m,\ \alpha KG}$ = 0.150 ± 0.003 mM; $k_{cat}/K_{m,\ \alpha KG\rightarrow D2HG}$ = 16.6 ± 0.4 $mM^{-1}$ $s^{-1}$. In all measurements, two biological replicates via protein preparations with each point showing the mean of the two biological replicates. Observed rate constants ($k_{obs}$) were calculated from the linear portion of plots of substrate concentration versus time. Kinetic parameters were calculated and reported as +/- SEM resulting from deviation of the mathematical fit.

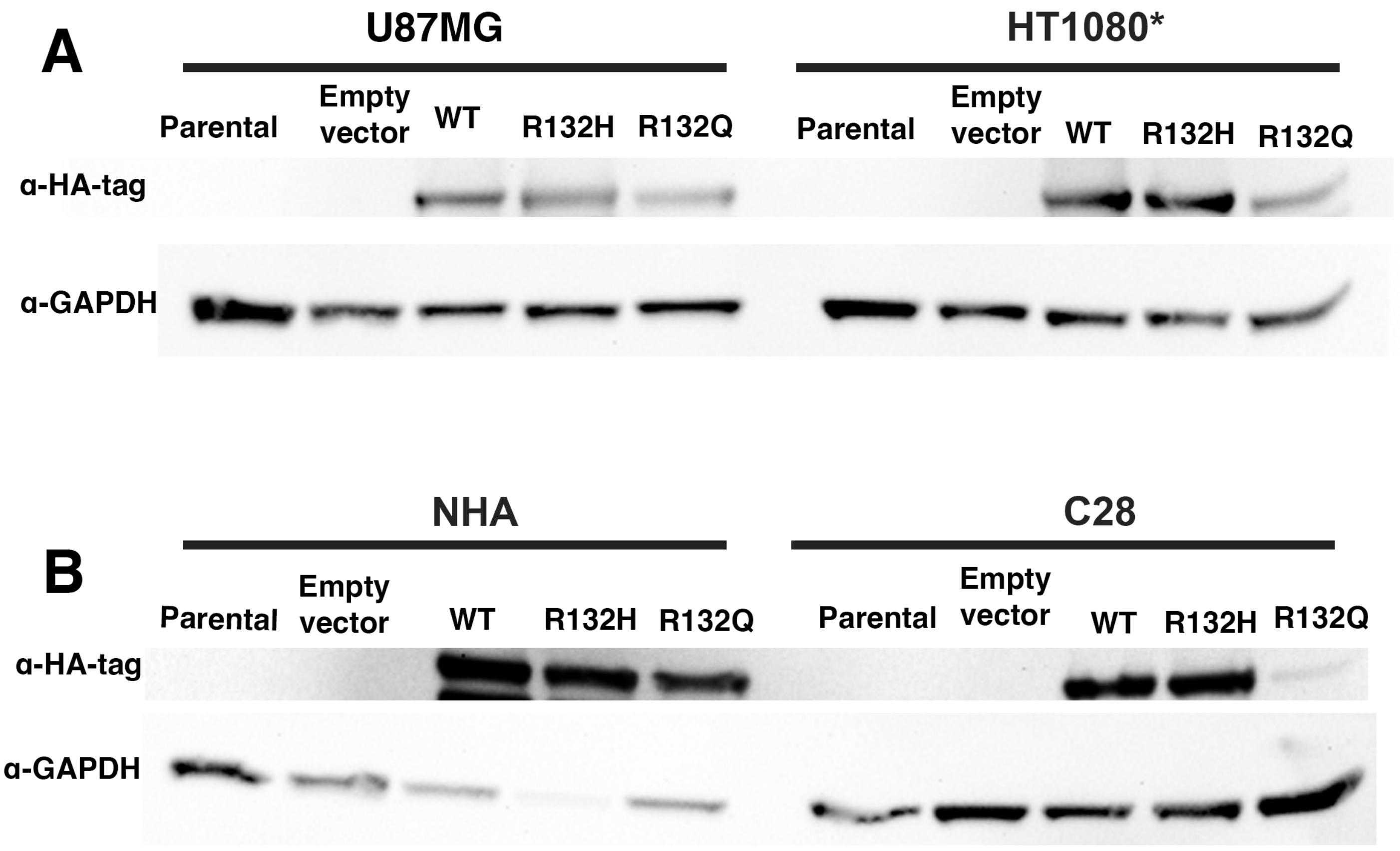


**Supplementary Fig. 2. Western immunoblot analysis of exogenously expressed HA-tagged IDH1. a**, U87MG and HT1080* cell lines stably overexpressing HA-tagged WT or mutant IDH1. **b**, Normal human astrocytes (NHA) and C28 cell lines stably overexpressing HA-tagged WT or mutant IDH1.

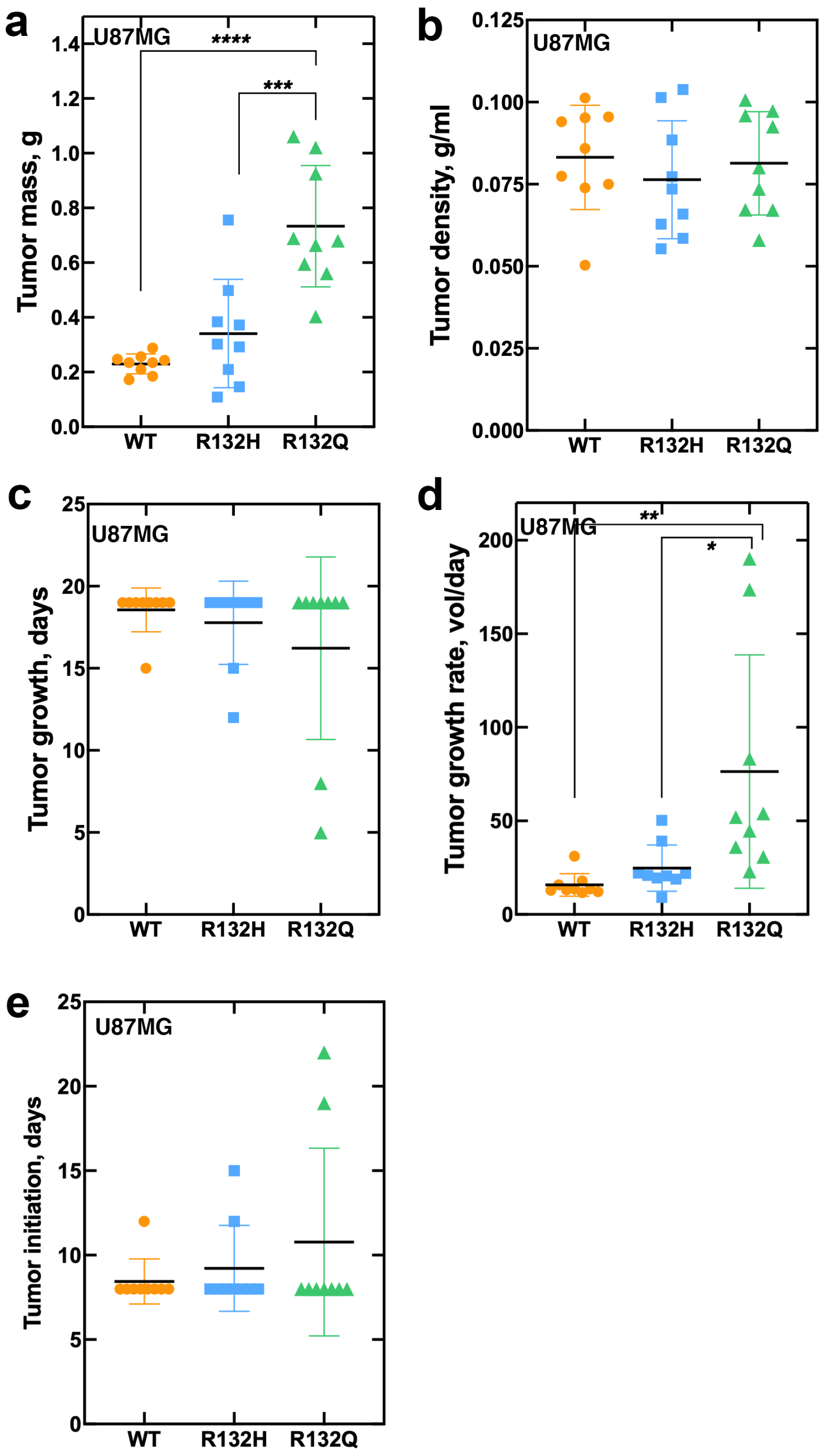


**Supplementary Fig. 3. Features of U87MG mouse xenografts.** Nine biological mouse xenograft replicates were generated from each of the U87MG cell lines stably overexpressing IDH1 WT (orange circles), R132H (blue squares), or R132Q (green triangles). In all panels, the median is shown as a bar and error bars are shown as mean ± SD. Ordinary one-way ANOVA analyses were performed, and if a significant change was determined, the p value is indicated ($p \leq 0.05$ (*), $p \leq 0.01$ (**), $p \leq 0.001$ (***), $p \leq 0.0001$ (****)). **a**, The mass of all tumors formed were weighed at the day of euthanasia. **b**, The density of each tumor was calculated as mass (in g) per volume density. **c**, Tumor growth days were calculated by determining the number of days from tumor injection day to euthanasia date, and subtracting the number of days it took for the tumor to initiate from this number. **d**, The tumor growth rate was calculated by dividing the number of tumor growth days by the tumor volume in mL using a 1 $mm^3$ = 0.01 mL conversion. **e**, Tumor initiation was determined by the number of days for a tumor to form after injection.

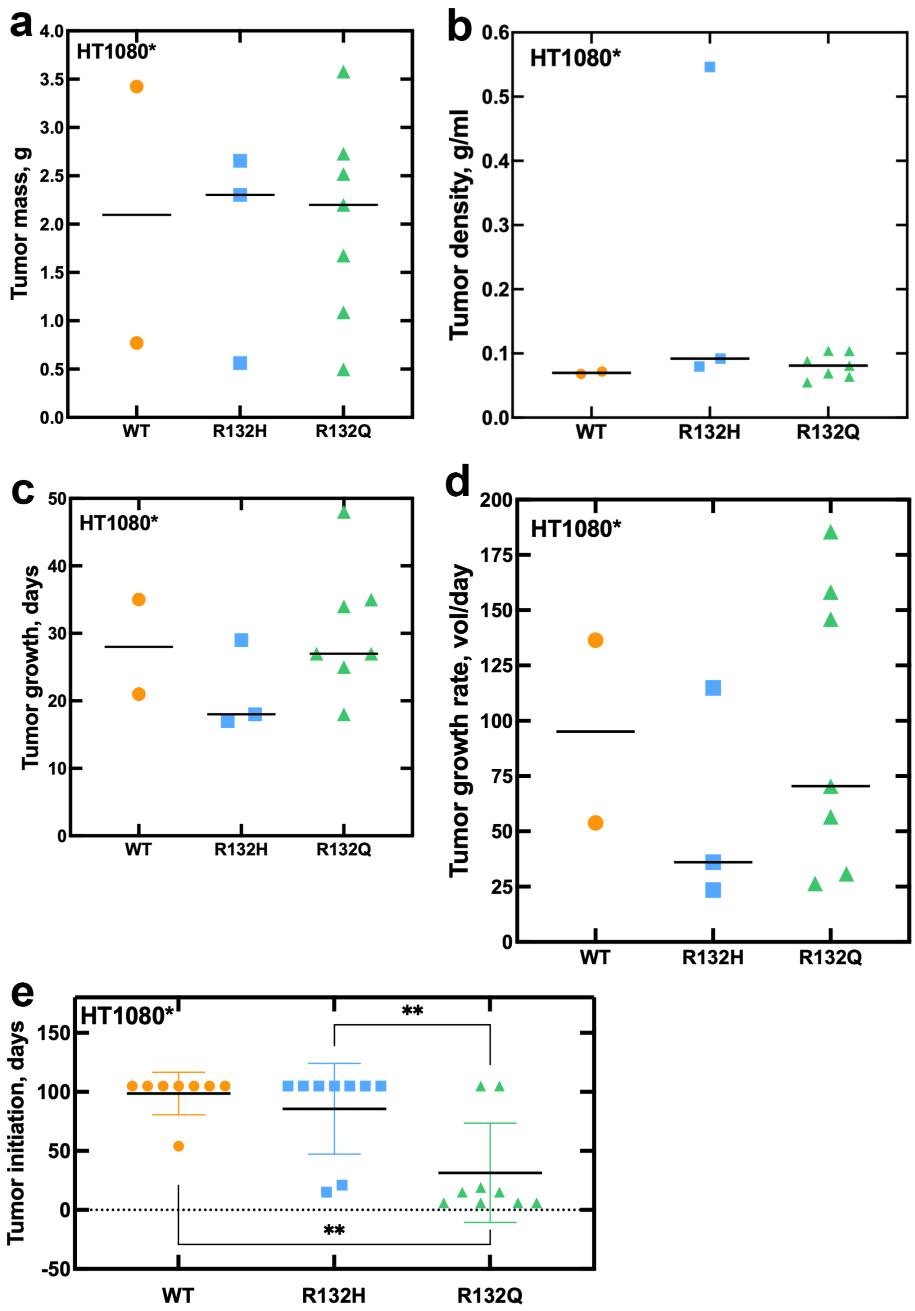


**Supplementary Fig. 4. Features of HT1080* mouse xenografts.** Nine biological mouse xenograft replicates were generated from each of the HT1080* cell lines stably overexpressing IDH1 WT (orange circles), R132H (blue squares), or R132Q (green triangles). However, not all biological replicates generated a tumor. In **a-d**, only biological replicates where tumors formed were analyzed. In **e**, biological replicates without tumors forming by day 105 are also included. In all panels, the median is shown as a bar. Ordinary one-way ANOVA analyses were performed, and if a significant change was determined, the p value is indicated ($p \leq 0.01$ (**)). **a**, The mass of all tumors formed were weighed at the day of euthanasia. **b**, The density of each tumor was calculated as mass (in g) per volume density. **c**, Tumor growth days were calculated by determining the number of days from tumor injection day to euthanasia date, and subtracting the number of days it took for the tumor to initiate from this number. **d**, The tumor growth rate was calculated by dividing the number of tumor growth days by the tumor volume in mL using a 1 $mm^3$ = 0.01 mL conversion. **e**, Tumor initiation was determined by the number of days for a tumor to form after injection. Here error bars are shown as mean ± standard deviation (S.D.).

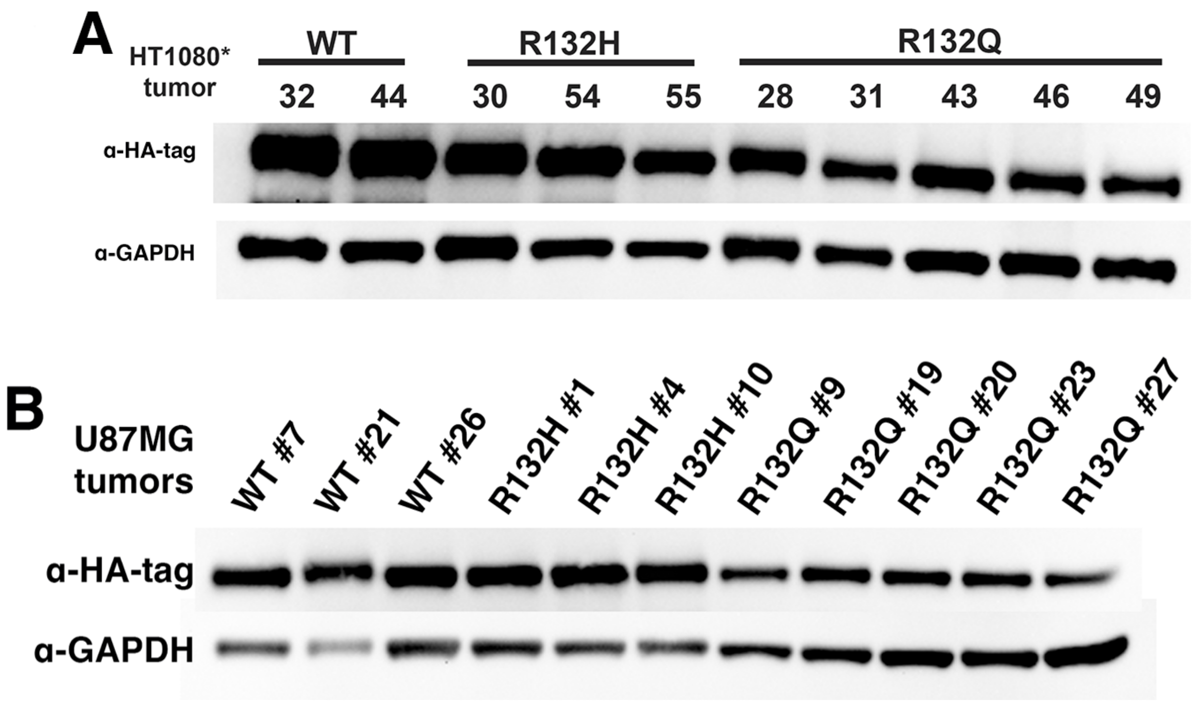


**Supplementary Fig. 5. Western immunoblot analysis of HT1080* and U87MG xenografts. a,** Western immunoblot analysis of HT1080* xenograft tumors expressing HA-tagged WT and mutant IDH1. **b**, Western immunoblot analysis of U87MG xenograft tumors expressing HA-tagged WT and mutant IDH1.

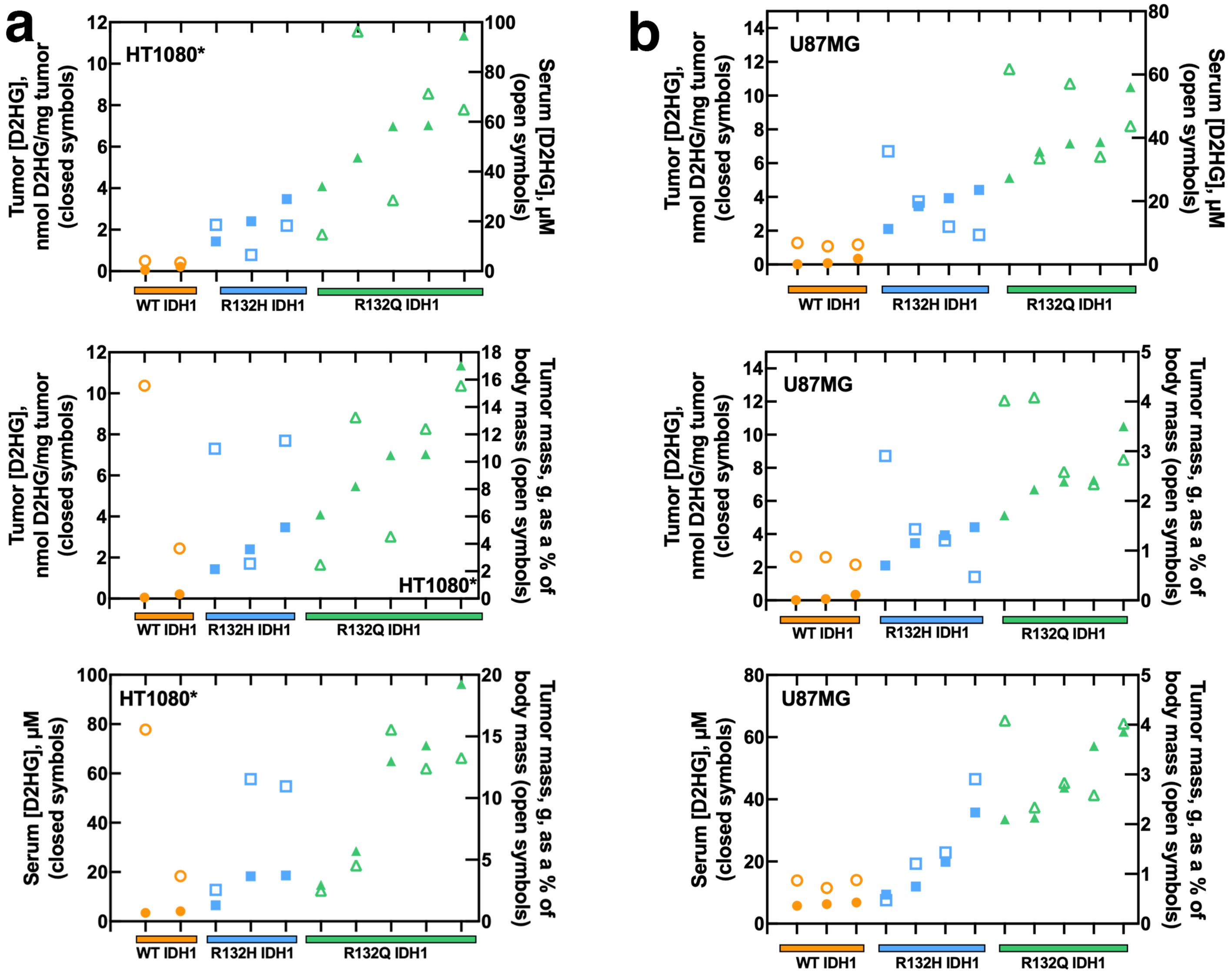


**Supplementary Fig. 6. Correlation of D2HG levels. a,** For each of the HT1080* xenograft tumors (2 biological replicates for WT, 3 biological replicates for R132H, and 5 biological replicates for R132Q), D2HG concentrations in tumors are plotted versus D2HG concentrations in serum (top) and versus tumor mass as a percent of body mass (middle). Serum D2HG concentrations are plotted versus tumor mass as a percent of body mass (bottom). **b,** For U87MG xenografts, D2HG concentrations in tumors are plotted versus D2HG concentrations in serum (top) and versus tumor mass as a percent of body mass (middle). Serum D2HG concentrations are plotted versus tumor mass as a percent of body mass (bottom). Each point represents a single biological replicate.

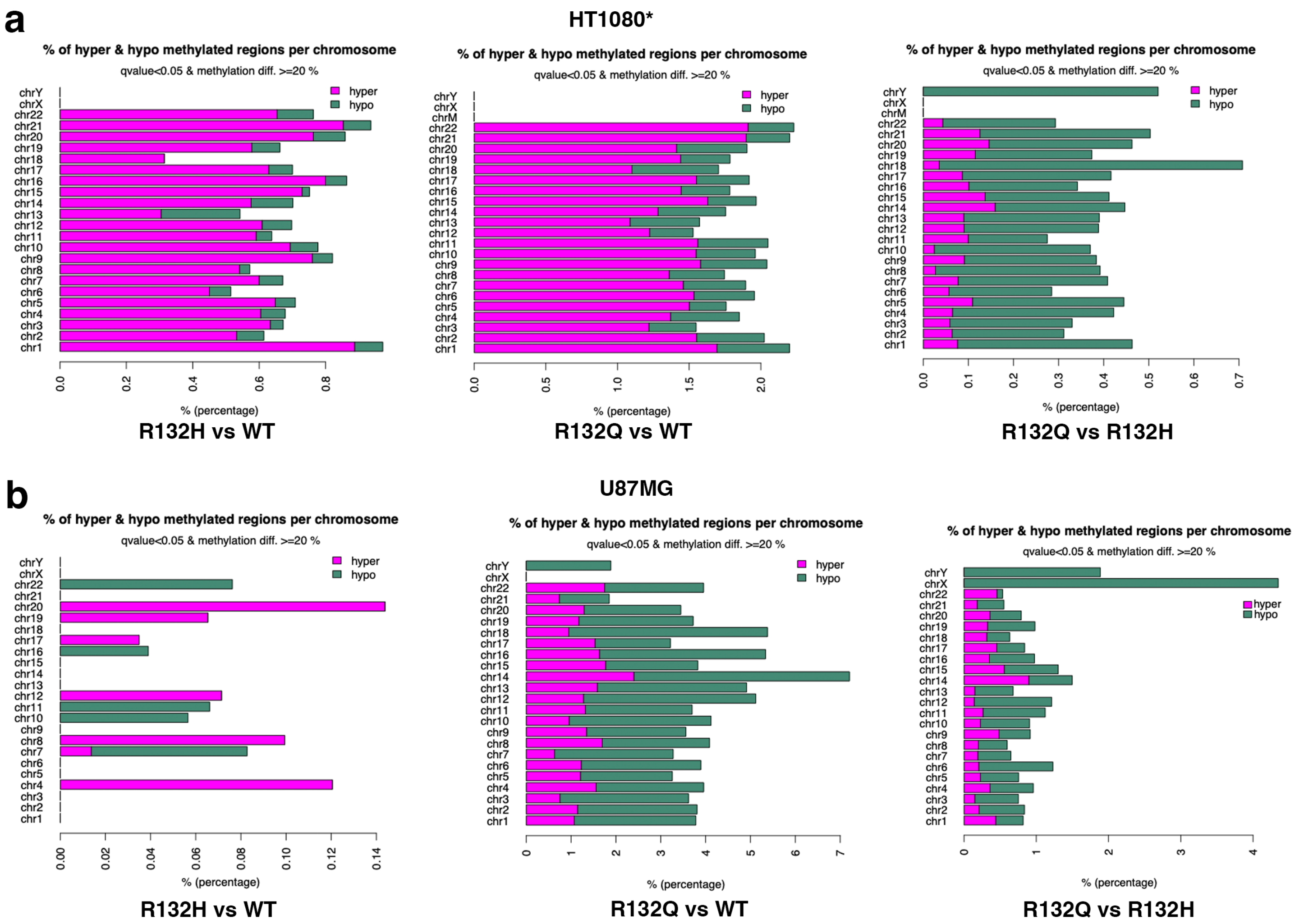


**Supplementary Fig. 7. Distribution of differentially methylated CpG sites.** The distribution of differentially methylated CpG sites (DMS, % methylation difference ≥ 20 and q-value < 0.05) per chromosome in **a**, HT1080* and **b**, U87MG tumor xenograft RRBS data. Left panels, DMS in IDH1 R132H compared to IDH1 WT; middle panels, DMS in IDH1 R132Q compared to IDH1 WT, right panels, DMS in IDH1 R132Q compared to IDH1 R132H tumor samples. Hypermethylated (hyper) DMS are shown in pink and hypomethylated (hypo) DMS are shown in green. Two of two, three of three, and five of seven (randomized selection) of the xenograft tumors that formed HT1080* xenograft tumors upon IDH1 WT, R132H, or R132Q expression, respectively, were evaluated as biological replicates by RRBS, though one of the R132H xenograft tumors failed analysis. Three of nine, four of nine, and five of nine (randomized selection for each) of the xenograft tumors that formed U87MG xenograft tumors upon IDH1 WT, R132H, or R132Q expression, respectively, were evaluated as biological replicates by RRBS.

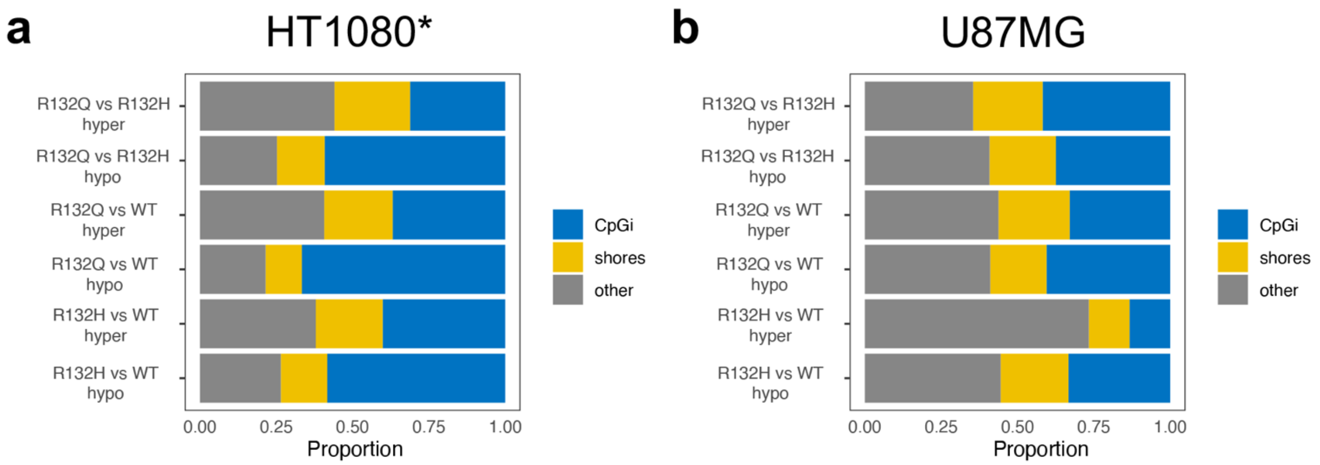


**Supplementary Fig. 8. Distribution of differentially methylated CpG sites.** Bar plots showing the proportion of the differentially methylated CpG sites by CpG island annotations in **a**, HT1080*, and **b**, U87MG tumor xenografts. Blue indicates CpG islands (CpGi), yellow indicates shores, and gray indicates other.

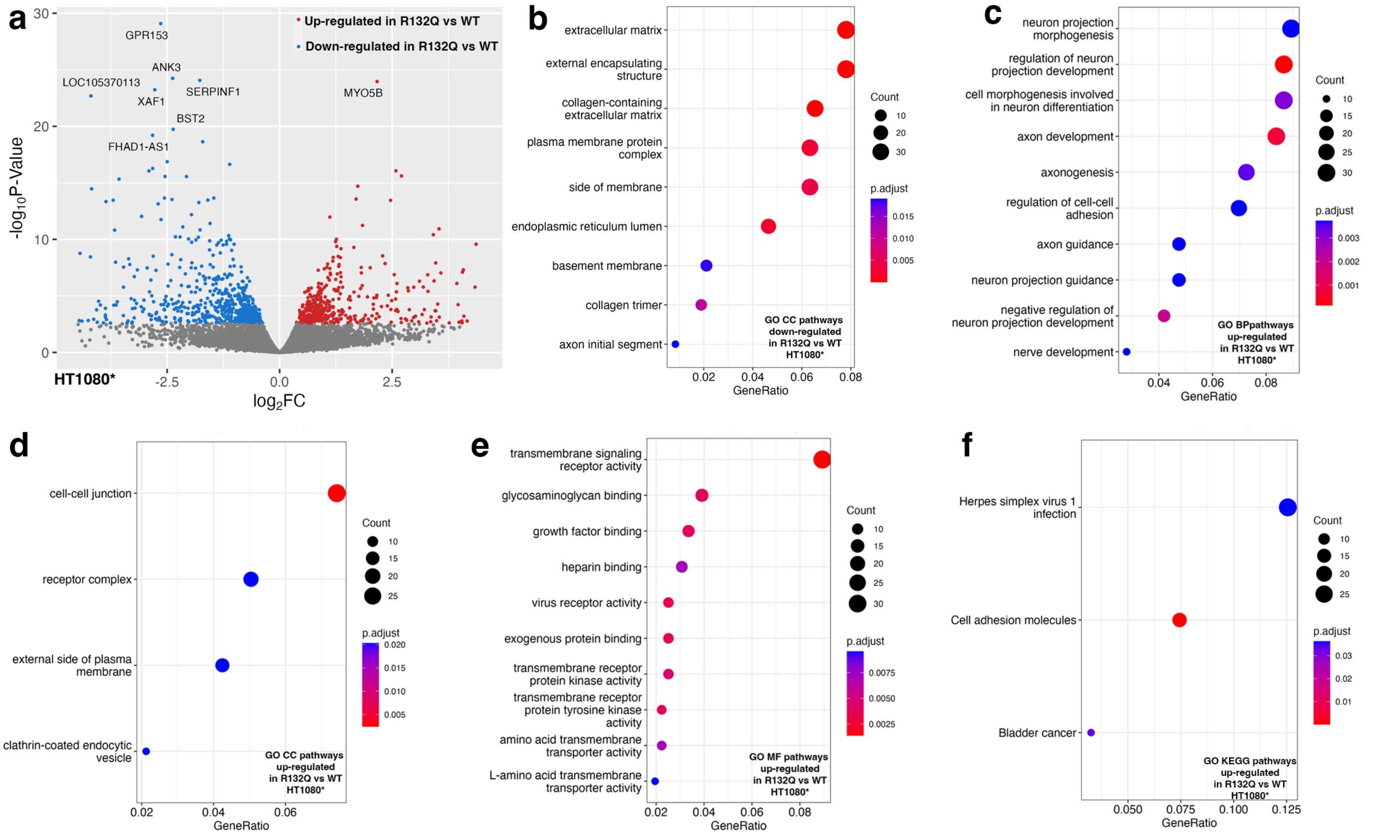


**Supplementary Fig. 9. Transcriptome analysis using RNAseq of HT1080* xenograft tumors comparing IDH1 R132Q and IDH1 WT**. Two of two, three of three, and five of seven (randomized selection) of the xenograft tumors that formed HT1080* xenograft tumors upon IDH1 WT, R132H, or R132Q expression, respectively, were evaluated as biological replicates by RNAseq. Down-regulated pathways refer to categories where most genes show decreased expression levels in mutant versus WT IDH1. Up-regulated pathways refer to categories where most genes show increased expression levels in mutant versus WT IDH1. Number of genes is indicated in the count, with the $p_{\text{adjusted}}$ value indicated by color. **a**, Volcano plot of differentially expressed transcripts comparing expression of IDH1 R132Q versus IDH1 WT. **b**, Cellular component (CC) pathways down-regulated in IDH1 R132Q versus IDH1 WT. **c**, Biological pathways (BP) up-regulated in IDH1 R132Q versus IDH1 WT. **d**, Cellular component (CC) pathways up-regulated in IDH1 R132Q versus IDH1 WT. **e**, Molecular function (MF) pathways up-regulated in IDH1 R132Q versus IDH1 WT. **f**, KEGG pathways up-regulated in IDH1 R132Q versus IDH1 WT.

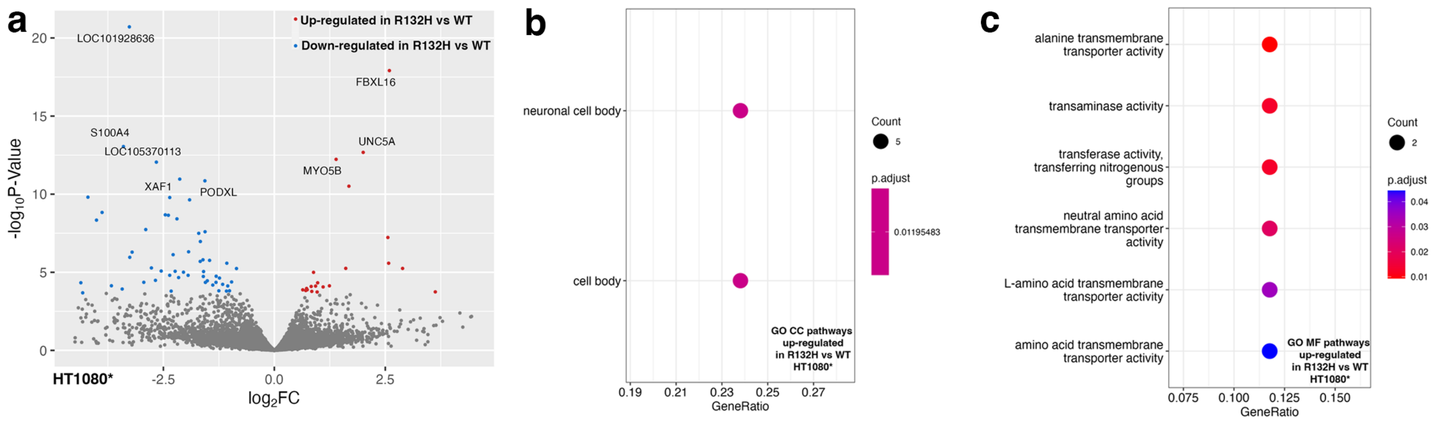


**Supplementary Fig. 10. Transcriptome analysis using RNAseq of HT1080* xenograft tumors comparing IDH1 R132H and IDH1 WT.** Two of two, three of three, and five of seven (randomized selection) of the xenograft tumors that formed HT1080* xenograft tumors upon IDH1 WT, R132H, or R132Q expression, respectively, were evaluated as biological replicates by RNAseq. Down-regulated pathways refer to categories where most genes show decreased expression levels in mutant versus WT IDH1. Up-regulated pathways refer to categories where most genes show increased expression levels in mutant versus WT IDH1. Number of genes is indicated in the count, with the $p_{\text{adjusted}}$ value indicated by color. **a**, Volcano plot of differentially expressed transcripts comparing expression of IDH1 R132H versus IDH1 WT. **b**, Cellular component (CC) pathways down-regulated in IDH1 R132H versus IDH1 WT. **c**, Molecular function (MF) pathways up-regulated in IDH1 R132H versus IDH1 WT.

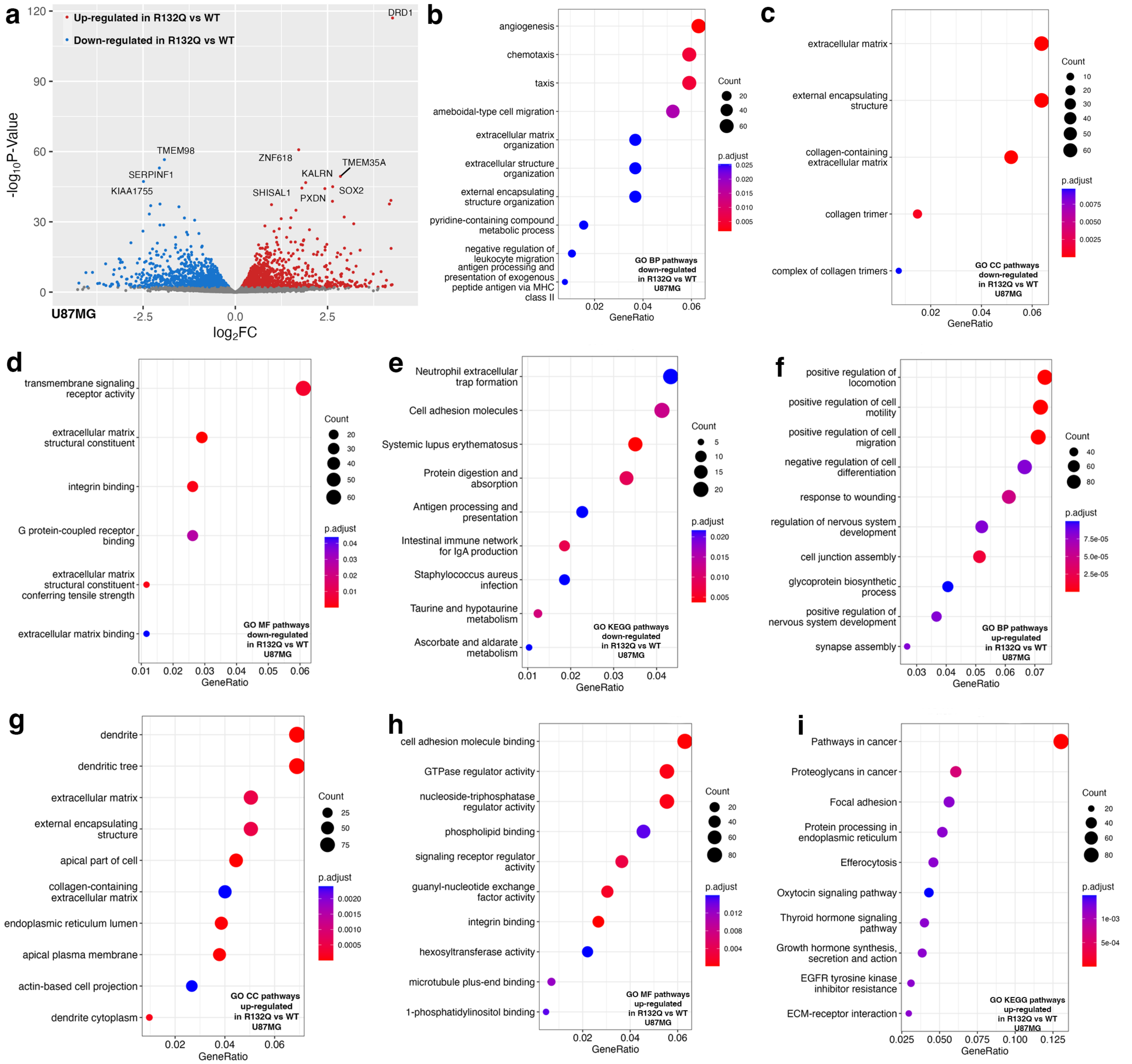


**Supplementary Fig. 11. Transcriptome analysis using RNAseq of U87MG xenograft tumors comparing IDH1 R132Q and IDH1 WT**. Three of nine, four of nine, and five of nine (randomized selection for each) of the xenograft tumors that formed U87MG xenograft tumors upon IDH1 WT, R132H, or R132Q expression, respectively, were evaluated as biological replicates by RNAseq. Down-regulated pathways refer to categories where most genes show decreased expression levels in mutant versus WT IDH1. Up-regulated pathways refer to categories where most genes show increased expression levels in mutant versus WT IDH1. Number of genes is indicated in the count, with the $p_{adjusted}$ value indicated by color. **a**, Volcano plot of differentially expressed transcripts comparing expression of IDH1 R132Q versus IDH1 WT. **b**, Biological pathways (BP) down-regulated in IDH1 R132Q versus IDH1 WT. **c**, Cellular component (CC) pathways down-regulated in IDH1 R132Q versus IDH1 WT. **d**, Molecular function (MF) pathways down-regulated in IDH1 R132Q versus IDH1 WT. **e**, KEGG pathways down-regulated in IDH1 R132Q versus IDH1 WT. **f**, BP up-regulated in IDH1 R132Q versus IDH1 WT. **g**, CC pathways up-regulated in IDH1 R132Q versus IDH1 WT. **h**, MF pathways up-regulated in IDH1 R132Q versus IDH1 WT. **i**, KEGG pathways up-regulated in IDH1 R132Q versus IDH1 WT.

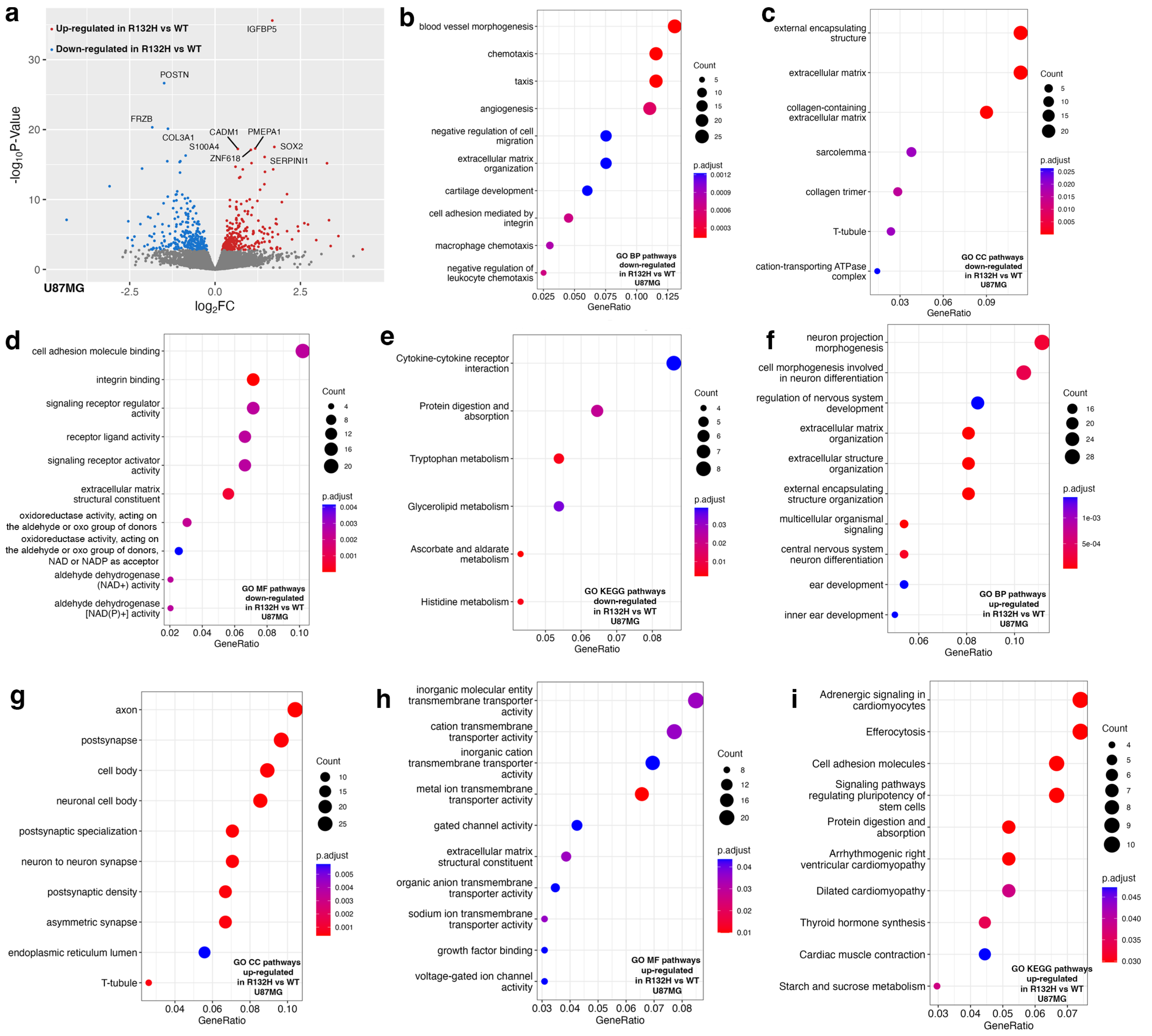

**Supplementary Fig. 12. Transcriptome analysis using RNAseq of U87MG xenograft tumors comparing IDH1 R132H and IDH1 WT**. Three of nine, four of nine, and five of nine (randomized selection for each) of the xenograft tumors that formed U87MG xenograft tumors upon IDH1 WT, R132H, or R132Q expression, respectively, were evaluated as biological replicates by RNAseq. Down-regulated pathways refer to categories where most genes show decreased expression levels in mutant versus WT IDH1. Up-regulated pathways refer to categories where most genes show increased expression levels in mutant versus WT IDH1. Number of genes is indicated in the count, with the $p_{\text{adjusted}}$ value indicated by color. **a**, Volcano plot of differentially expressed transcripts comparing expression of IDH1 R132H versus IDH1 WT. **b**, Biological pathways (BP) down-regulated in IDH1 R132H versus IDH1 WT. **c**, Cellular component (CC) pathways down-regulated in IDH1 R132H versus IDH1 WT. **d**, Molecular function (MF) pathways down-regulated in IDH1 R132H versus IDH1 WT. **e**, KEGG pathways down-regulated in IDH1 R132H versus IDH1 WT. **f**, BP up-regulated in IDH1 R132H versus IDH1 WT. **g**, CC pathways up-regulated in IDH1 R132H versus IDH1 WT. **h**, MF pathways up-regulated in IDH1 R132H versus IDH1 WT. **i**, KEGG pathways up-regulated in IDH1 R132H versus IDH1 WT.

**Supplementary Table 1.** Features of HT1080* and U87MG tumor xenografts.

| Xenograft | IDH1 (ID #) | Tumor weight, g, as a % of body weight | [Tumor 2HG], nmol tumor D2HG/mg tumor | [Serum 2HG], μM | Number of covered CpGs (minCov > 10) | IDH1 transcripts (RNAseq) |
|---|---|---|---|---|---|---|
| HT1080* | WT (32) | 3.67 | 0.21 | 4.12 | 1,688,753 | 26,331 |
| HT1080* | WT (44) | 15.56 | 0.057 | 3.46 | 2,098,845 | 41,577 |
| HT1080* | R132H (30) | 10.96 | 1.44 | 18.63 | 1,548,982 | 11,325 |
| HT1080* | R132H (54) | 11.55 | 3.48 | 18.33 | 2,102,976 | 28,180 |
| HT1080* | R132H (55) | 2.56 | 2.40 | 6.50 | RRBS analysis failed | 10,981 |
| HT1080* | R132Q (28) | 2.49 | 4.10 | 14.83 | 1,056,926 | 9,501 |
| HT1080* | R132Q (31) | 4.54 | 6.99 | 28.56 | 1,815,817 | 12,003 |
| HT1080* | R132Q (34) | 10.0 | 1.14 | 29.74 | ND | ND |
| HT1080* | R132Q (43) | 12.41 | 7.04 | 71.40 | 1,996,037 | 13,976 |
| HT1080* | R132Q (46) | 15.56 | 11.36 | 64.94 | 2,247,374 | 9,553 |
| HT1080* | R132Q (49) | 13.25 | 5.48 | 96.28 | 1,616,142 | 12,640 |
| HT1080* | R132Q (52) | 7.61 | 4.67 | 45.60 | ND | ND |
| U87MG | WT (7) | 0.88 | 0.021 | 6.81 | 1,686,541 | 18,566 |
| U87MG | WT (18) | 1.07 | 0.046 | 6.23 | ND | ND |
| U87MG | WT (11) | 1.07 | 0.064 | 5.31 | ND | ND |
| U87MG | WT (13) | 0.80 | 0.073 | 6.65 | ND | ND |
| U87MG | WT (21) | 0.87 | 0.075 | 5.76 | 1,670,198 | 18,608 |
| U87MG | WT (17) | 1.01 | 0.10 | 5.67 | ND | ND |
| U87MG | WT (2) | 1.07 | 0.14 | 10.20 | ND | ND |
| U87MG | WT (26) | 0.72 | 0.34 | 6.27 | 1,643,486 | 15,668 |
| U87MG | WT (6) | 1.25 | 0.55 | 6.19 | ND | ND |
| U87MG | R132H (1) | 2.91 | 2.11 | 35.78 | 1,889,953 | 20,244 |
| U87MG | R132H (15) | 1.74 | 2.77 | 17.55 | ND | ND |
| U87MG | R132H (16) | 0.91 | 3.29 | 9.54 | ND | ND |
| U87MG | R132H (4) | 1.43 | 3.45 | 19.89 | 1,794,986 | 16,962 |
| U87MG | R132H (25) | 0.63 | 3.47 | 11.60 | ND | ND |
| U87MG | R132H (12) | 2.26 | 3.48 | 15.83 | RRBS analysis failed | ND |
| U87MG | R132H (22) | 1.32 | 3.63 | 14.89 | ND | ND |
| U87MG | R132H (10) | 1.21 | 3.94 | 11.97 | 1,858,544 | 21,612 |
| U87MG | R132H (24) | 0.47 | 4.42 | 9.39 | 1,763,802 | 18,808 |
| U87MG | R132Q (20) | 4.02 | 5.13 | 61.76 | 1,395,526 | 12,249 |
| U87MG | R132Q (3) | 2.77 | 5.86 | 54.93 | ND | ND |
| U87MG | R132Q (5) | 4.42 | 6.04 | 42.28 | ND | ND |
| U87MG | R132Q (14) | 2.87 | 6.07 | 47.51 | ND | ND |
| U87MG | R132Q (9) | 4.08 | 6.70 | 33.58 | 1,841,212 | 12,188 |
| U87MG | R132Q (27) | 2.59 | 7.17 | 57.15 | 1,531,687 | 12,688 |
| U87MG | R132Q (8) | 1.75 | 7.22 | 34.46 | ND | ND |
| U87MG | R132Q (19) | 2.34 | 7.25 | 34.07 | 782,672 | 10,521 |
| U87MG | R132Q (23) | 2.83 | 10.51 | 43.77 | 1,567,435 | 11,596 |

**Supplementary Table 2.** Differentially expressed genes in pairwise comparisons of RNA-seq analysis.

| | **R132H vs WT HT1080*** | **R132Q vs WT HT1080*** | **R132Q vs WT HT1080*** | **R132H vs WT U87MG** | **R132Q vs WT U87MG** | **R132H vs R132Q U87MG** |
|---|---|---|---|---|---|---|
| Total number of genes | 20002 | 20642 | 18643 | 18754 | 21076 | 17868 |
| Total number of significant genes (padj < 0.05) | 83 | 1062 | 959 | 573 | 2994 | 801 |
| Total number of significant genes (padj < 0.05 and abs(log2FC) > 1) | 71 | 592 | 327 | 153 | 837 | 173 |
| Total number of significantly upregulated genes (padj < 0.05 and abs(log2FC) > 0) | 21 | 470 | 518 | 307 | 1619 | 439 |
| Total number of significantly upregulated genes (padj < 0.05 and abs(log2FC) > 1) | 11 | 247 | 257 | 68 | 389 | 81 |
| Total number of significantly downregulated genes (padj < 0.05 and abs(log2FC) > 0) | 62 | 592 | 441 | 266 | 1375 | 362 |
| Total number of significantly downregulated genes (padj < 0.05 and abs(log2FC) > 1) | 60 | 345 | 70 | 85 | 448 | 92 |

**Supplementary Table 3.** RNAseq transcript details for selected genes from HT1080* tumor xenografts. The relative change in Log2 FC is shown in a blue (relative decrease in transcripts) to red (relative increase of transcripts), and significantly altered genes based on $p_{adj}$ values are notated in a white (non-significant) to grey (increasing significance) scale.

| | R132Q vs R132H | | R132Q vs WT | | R132H vs WT | |
|---|---|---|---|---|---|---|
| GeneID | Log2 FC | padj | Log2 FC | padj | Log2 FC | padj |
| AFDN | 0.52 | 0.00 | 0.68 | 0.00 | 0.15 | 1.00 |
| ANXA6 | -0.38 | 0.02 | -0.48 | 0.03 | -0.09 | 1.00 |
| CBL | 0.45 | 0.02 | 0.46 | 0.25 | 0.01 | 1.00 |
| CASP9 | -0.65 | 0.00 | -0.48 | 0.30 | 0.18 | 1.00 |
| COL9A2 | 0.49 | 0.78 | -2.52 | 0.04 | -3.01 | 0.28 |
| CCND2 | 3.83 | 0.04 | 2.22 | 0.39 | NA | NA |
| CDK6 | 0.70 | 0.00 | 0.50 | 0.17 | -0.19 | 1.00 |
| EGFR | 0.99 | 0.00 | 1.10 | 0.00 | 0.12 | 1.00 |
| ETS1 | 0.67 | 0.00 | 0.77 | 0.01 | 0.11 | 1.00 |
| FZD1 | -0.65 | 0.00 | -0.72 | 0.11 | -0.07 | 0.91 |
| FZD9 | NA | NA | -1.94 | 0.14 | -2.50 | 0.71 |
| JAG1 | 1.15 | 0.00 | 1.01 | 0.01 | -0.14 | 1.00 |
| NKD1 | -0.30 | 0.72 | -1.41 | 0.00 | -1.10 | 0.92 |
| MYC | 0.94 | 0.00 | 1.31 | 0.00 | 0.37 | 1.00 |
| RALA | 0.52 | 0.00 | 0.32 | 0.55 | -0.20 | 1.00 |
| RASAL2 | 0.72 | 0.00 | 0.72 | 0.00 | 0.00 | 1.00 |
| TCF7 | 0.50 | 0.76 | -0.59 | 0.80 | -1.09 | 1.00 |
| TP53 | -0.12 | 0.68 | -0.23 | 0.44 | -0.10 | 1.00 |
| TP53I3 | -0.40 | 0.03 | -0.79 | 0.00 | -0.38 | 1.00 |
| VEGFC | 0.81 | 0.00 | 0.48 | 0.14 | -0.32 | 1.00 |
| WNT9A | -0.31 | 0.41 | -1.07 | 0.00 | -0.75 | 0.52 |
| WNT10B | 0.95 | 0.06 | 2.09 | 0.00 | 1.15 | 1.00 |

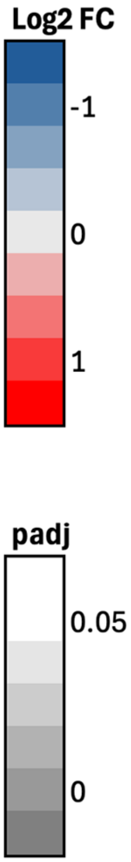

**Supplementary Table 4.** RNAseq transcript details for selected genes from U87MG tumor xenografts. The relative change in Log2 FC is shown in a blue (relative decrease in transcripts) to red (relative increase of transcripts), and significantly altered genes based on $p_{adj}$ values are notated in a white (non-significant) to grey (increasing significance) scale.

| | R132Q vs R132H | | R132Q vs WT | | R132H vs WT | |
|---|---|---|---|---|---|---|
| **GeneID** | Log2 FC | padj | Log2 FC | padj | Log2 FC | padj |
| AFDN | 0.26 | 0.02 | 0.24 | 0.01 | -0.02 | 0.98 |
| CBL | 0.32 | 0.05 | 0.39 | 0.00 | 0.07 | 0.94 |
| CDK6 | 0.18 | 0.47 | 0.30 | 0.08 | 0.12 | 0.77 |
| COL4A5 | 0.61 | 0.01 | 1.16 | 0.00 | 0.55 | 0.00 |
| COL4A6 | 0.82 | 0.00 | 1.62 | 0.00 | 0.80 | 0.00 |
| EGFR | 0.56 | 0.00 | 0.62 | 0.00 | 0.05 | 0.95 |
| ETS1 | 0.35 | 0.02 | 0.40 | 0.00 | 0.05 | 0.95 |
| FZD1 | -0.47 | 0.00 | -0.61 | 0.00 | -0.14 | 0.75 |
| ITGA2 | 0.99 | 0.00 | 1.49 | 0.00 | 0.50 | 0.00 |
| ITGAV | 0.49 | 0.00 | 0.61 | 0.00 | 0.12 | 0.69 |
| ITGB1 | 0.27 | 0.05 | 0.48 | 0.00 | 0.22 | 0.21 |
| IL6 | 3.24 | 0.05 | 259.00 | 0.04 | 0.07 | 0.99 |
| JAG1 | 0.40 | 0.04 | 0.78 | 0.00 | 0.38 | 0.07 |
| JAK1 | 0.22 | 0.01 | 0.26 | 0.01 | 0.03 | 0.97 |
| JUN | 0.37 | 0.03 | 0.60 | 0.00 | 0.23 | 0.40 |
| MET | 0.30 | 0.10 | 0.48 | 0.00 | 0.17 | 0.60 |
| MDM2 | 0.35 | 0.03 | 0.19 | 0.35 | -0.16 | 0.53 |
| NRAS | 0.19 | 0.12 | 0.26 | 0.01 | 0.06 | 0.90 |
| RASAL2 | 0.27 | 0.07 | 0.37 | 0.00 | 0.09 | 0.88 |
| ROCK2 | 0.32 | 0.00 | 0.23 | 0.03 | -0.09 | 0.83 |
| STAT3 | 0.23 | 0.08 | 0.44 | 0.00 | 0.20 | 0.12 |
| TGFA | 0.64 | 0.47 | 1.36 | 0.01 | 0.72 | 0.16 |
| TGFB1 | 0.05 | 0.89 | 0.36 | 0.00 | 0.31 | 0.07 |
| TP53 | -0.22 | 0.42 | -0.37 | 0.03 | -0.15 | 0.69 |
| TP53I3 | -0.16 | 0.67 | -0.55 | 0.00 | -0.40 | 0.11 |
| WNT10B | -1.26 | 0.00 | -1.70 | 0.00 | -0.44 | 0.67 |

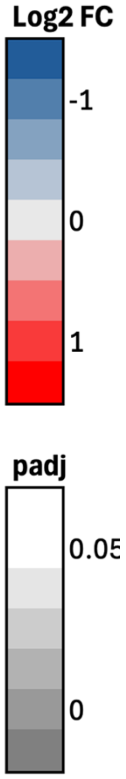
